# A global perspective on the genomics of Moraxella catarrhalis

**DOI:** 10.1101/2025.03.28.645966

**Authors:** Makarena Gonzalez-Reyes, Ignacio Ramos-Tapia, Juan A. Ugalde

## Abstract

2.

*Moraxella catarrhalis* is an opportunistic pathogen of the human respiratory tract, primarily associated with otitis media in children and exacerbations of chronic obstructive pulmonary disease (COPD) in adults. Despite its clinical importance, the genomic diversity and functional specialization of *M. catarrhalis* remain insufficiently characterized. This study aimed to analyze the global genetic diversity of *M. catarrhalis* using whole-genome sequencing to identify phylogenetic lineages, antimicrobial resistance patterns, and key virulence factors. Phylogenomic analysis of 345 publicly available genomes identified three phylogroups, of which one exhibited significant genomic divergence and was excluded from further analyses due to its potential classification as a separate species. The remaining two phylogroups corresponded to previously described seroresistant and serosensitive lineages. Phylogroup B exhibited a higher prevalence of antimicrobial resistance genes, particularly *bro-1* and *bro-2*, while Phylogroup A exhibited unique metabolic adaptation, including genes encoding for the DppB-DppC-DppD dipeptide transport system. Both phylogroups shared crucial virulence factors, including UspA1 and UspA2, which facilitate adhesion and immune evasion. Potential therapeutic targets were identified, including PilQ, essential for type IV pilus biogenesis, and CopB, which plays a key role in iron acquisition and immune evasion. Overall, these findings highlight the significance of phylogenomics approaches to elucidate the genetic mechanisms underlying pathogenicity and resistance in *M. catarrhalis*, providing insights for future therapeutic and preventive strategies.

**3. Impact statement:** This study presents a comprehensive phylogenomic analysis of *M. catarrhalis*, expanding current knowledge of its genetic diversity, antimicrobial resistance, and virulence factors. By analyzing 345 global genomes, we identified three distinct phylogroups, one of which shows substantial genomic divergence, suggesting it may represent a novel species within the *Moraxella* genus. Our findings confirm the existence of seroresistant and serosensitive lineages, providing new insights into their metabolic adaptations and differential antibiotic resistance profiles. Notably, phylogroup B harbors a higher prevalence of β-lactam resistance genes, while phylogroup A exhibits unique peptide transport systems. This work strengthens the evidence that phylogenomic approaches are essential to understand the evolutionary dynamics and pathogenic potential of *M. catarrhalis*. The identification of conserved virulence factors and potential therapeutic targets, such as PilQ and CopB, underscores the relevance of this pathogen in respiratory infections and the need for targeted interventions. Our results not only clarify the population structure of *M. catarrhalis* but also provide a valuable genomic framework for future studies on antimicrobial resistance and vaccine development, with broad utility for microbiologists, clinicians, and public health researchers.

**4. Data summary:** The authors confirm all supporting data, code, and protocols have been provided within the article or through supplementary data files.

## 5. Introduction

*Moraxella catarrhalis* is a Gram-negative bacteria from the order Pseudomonadales, recognized as an opportunistic pathogen implicated in both upper and lower respiratory tract infections in humans [1]. Its prevalence is associated with respiratory tract infections and exhibits notable seasonal variation, with a higher incidence in winter [2]. In the upper respiratory tract, this bacteria is one of the leading causes of otitis media, alongside *Streptococcus pneumoniae* and *Haemophilus influenzae* [3,4], accounting for 15% to 20% of acute otitis media episodes in children [5–7]. Additionally, it is responsible for approximately 20% of acute bacterial sinusitis cases in children and a smaller proportion in adults [7–9]. In the lower respiratory tract, *M. catarrhalis* is a major etiological agent of infections such as pneumonia and exacerbations of chronic obstructive pulmonary disease (COPD) in adults, causing an estimated 2–4 million cases annually in the United States, which accounts for about 10% of all exacerbations [7,10]. In immunocompromised individuals, *M. catarrhalis* can lead to severe infections, including septicemia, meningitis, and endocarditis [11–14]. Furthermore, hospital outbreaks of respiratory diseases linked to this microorganism have been reported [14–16], highlighting its clinical and epidemiological importance in both community and healthcare settings.

Previous studies using molecular typing methods, including multilocus sequence typing (MLST), 16S and 23S rRNA gene sequencing, fingerprinting of outer membrane proteins, and restriction fragment polymorphism (RFLP) [17–21] have suggested that *M. cattarhalis* is composed of two distinct lineages, seroresistant (SR) and serosensitive (SS), with different virulence potential [19]. In addition, divergent strains with limited genetic homology to these lineages have also been described [21]. The SR lineage, primarily associated with 16S ribotype (RB) 1 strains [17,19], exhibits a highly pathogenic profile due to two key virulence traits: high resistance to the human complement system and efficient adhesion to respiratory epithelial cells [21]. This lineage is more frequently associated with respiratory tract diseases, as 51% of isolates are derived from diseased individuals [19,22]. In contrast, the SS lineage, composed of strains with RB2 and RB3 ribotypes [17,19], is sensitive to complement-mediated killing and demonstrates lower adhesion efficiency, being less commonly linked to disease (15% of isolates from diseased individuals) [19,22]. Despite these differences, both lineages retain a conserved set of essential genes involved in fundamental cellular processes, supporting their classification within the same species. However, their independent evolutionary trajectories indicate that they diverged from a common ancestor and evolved separately [19], reflecting their genetic and adaptive complexity.

*M. catarrhalis* has developed several strategies to evade antibiotic action, notably through three primary mechanisms: production of β-lactamases, efflux pumps, and biofilm formation. The main resistance strategy employed by *M. catarrhalis* involves the production of β-lactamase enzymes [23], specifically the *BRO-1*and *BRO-2* variants, which can inactivate penicillin and other β-lactam antibiotics [24]. Currently, more than 95% of clinical isolates exhibit resistance to penicillin [25,26]. In addition to β-lactamase production, *M. catarrhalis* possesses resistance-nodulation-division (RND)-type efflux pumps, specifically AcrAB-OprM, which confer intrinsic resistance against multiple classes of antimicrobials, including β-lactams, quinolones, and aminoglycosides [27,28]. These resistance mechanisms can be activated by environmental stress, such as a cold shock at 26°C, or by exposure to antibiotics themselves, thus enhancing antibiotic resistance through active expulsion of the compounds [28,29]. Another critical strategy used by *M. catarrhalis* to resist antibiotic action is biofilm formation. Biofilms are bacterial communities adhered to surfaces and embedded in an extracellular matrix that acts as a physical barrier against antimicrobial agents [30,31]. Biofilm formation is directly associated with persistent infections, such as chronic otitis media, where *M. catarrhalis* has been detected alongside other respiratory pathogens within biofilms [30]. Bacteria within these biofilms exhibit altered metabolism and reduced susceptibility to antibiotics, while maintaining conventional resistance mechanisms such as β-lactamase production [32,33].

Among the factors contributing to the pathogenesis of *M. catarrhalis*, the lipooligosaccharide (LOS) and several outer membrane proteins (OMPs) play a pivotal role [34,35], with the ubiquitous surface protein A (UspA) standing out as a key element [35]. UspA is a high-molecular-weight, multifunctional trimeric adhesin abundantly expressed on the bacterial surface [36], where it facilitates adherence and evasion of the host immune system [37]. In the *M. catarrhalis* population, three genes encoding UspA variants; *uspA1*, *uspA2*, and *uspA2H* have been identified. Clinical studies indicate that *uspA1* is present in 99% of isolates [38], while *uspA2* or *uspA2H* are expressed in a ratio of approximately 4:1 [38,39]. In terms of pathogenicity, UspA1 and UspA2H are primarily associated with biofilm formation and adherence to respiratory tract epithelial cells [34,40,41], facilitating bacterial colonization. On the other hand, UspA2 and, in some cases, UspA2H, contribute to serum resistance by binding host complement inhibitors, such as vitronectin and C4b-binding protein (C4BP) [42,43].

The clinical and epidemiological relevance of *M. catarrhalis* lies in its remarkable adaptability, antimicrobial resistance, and genetic diversification. However, despite advances in its genomic characterization, a clear genetic basis explaining the differences in virulence among its lineages has not been identified. Studies conducted so far have revealed significant evolutionary complexity but have failed to establish consistent links between lineages, virulence factors, and the metadata associated with each strain, such as the year of collection, clinical presentation, and geographic location. To address these limitations, it is essential to employ a genomic model that captures the genetic diversity of the species. This approach allows for the exploration of how the genetic variability of *M. catarrhalis* influences virulence, antimicrobial resistance, and other phenotypic traits. Using publicly available genomic data, this study aims to correlate phylogenetic lineages with key genomic characteristics, providing new perspectives to understand the determinants of pathogenicity in this species.

## 6. Methodssoftware mlst v.23.0

### 6.1 *M. catarrhalis* genomes acquisition

A total of 217 public genomes [44] and 3,135 Sequence Read Archive (SRA) [45] files were retrieved from the National Center for Biotechnology Information (NCBI). The search was conducted using the term "Moraxella catarrhalis" and the data were retrieved on March 30, 2024. The SRA reads were cleaned using fastp v0.23.4 [46,47], removing low-quality sequences and adapters. Subsequently, The reads were evaluated to determine if they correspond to *M. catarrhalis* using Kraken2 v2.1.3 [48,49], with the Standard database, which includes bacteria, archaea, human, plasmid, UniVec_Core, and viral, applying a selection threshold ≥98% for the percentage of fragments covered by the clade rooted in a taxon, ensuring a reliable identification of the microorganism. To determine whether any reads matched previously downloaded genomes, alignments were performed between the reads and the genomes. A read was considered distinct from the assembled genomes if the percentage of mapped reads and properly paired reads was ≤98%, and the number of singleton reads (unpaired reads) was ≥ 0.02%. The reads were assembled using *SPAdes* v3.15.5 [50], and the quality of both the assemblies and the previously downloaded genomes was verified using *CheckM2* v1.0.1 [51], ensuring a completeness ≥98% and a contamination level ≤2%, according to the criteria published by Chklovski et al. [51] subsequently, multilocus sequence typing was performed using the software mlst v.23.0 [52–54] was performed to characterize the strains by assigning alleles to the loci *abcZ*, *adk*, *efp*, *fumC*, *glyBeta*, *mutY*, *ppa*, and *trpE* [55]. After applying the filters, a total of 345 genomes were selected for further analysis. For more details on the applied filters, see Table S1.

### 6.2 Phylogenomic analysis

Gene prediction and annotation was performed using Bakta v1.8.1 [56]. The files generated in this process served as the basis for the pangenome analysis conducted with Panaroo v1.3.3 [57], applying a 100% threshold for the core genome. Subsequently, the core genomes were aligned using a base substitution model (GTR) combined with a categorical approximation (CAT) to handle the heterogeneity in evolutionary rates across different sites in the alignment (GTRCAT), and phylogenetic reconstruction was performed using maximum likelihood with RAxML v8.2.12 [58–61]. To validate the robustness of the phylogenetic relationships, support values were calculated from 1,000 bootstrap replicates. Phylogenetic clustering was carried out using TreeCluster v1.0.4 [62], applying the single-linkage clustering method, which groups clusters based on the shortest pairwise distance between elements of different clusters [63], with a branch length threshold of 0.002.

### 6.3 Pangenome analysis

Based on the gene presence-absence matrix generated by Panaroo v1.3.3 [57], the genes were classified into two main categories: core genes, present in 100% of the strains, and accessory genes, which are further subdivided into Soft core genes (95% ≤ strains < 100%), shell genes (15% ≤ strains < 95%), and cloud genes (strains < 15%) [64]. Subsequently, using the same matrix, the Heap’s Law was calculated [65,66], which describes the relationship between the number of genomes analyzed and the number of unique genes identified in the pangenome, determining the growth exponent (γ), a value that is used to classify the pangenome as closed if γ < 0, open if γ > 0, or reaching a fixed size (neither open nor closed) if γ = 0 [65].

### 6.4 Functional Ontology Assignments

For functional ontology assignment, STRING v12.0 was used, which implements Gene Ontology (GO) as a classification system for gene-set enrichment analysis [67]. The analysis was performed using the amino acid sequences obtained from Bakta v1.8.1 [56], considering *M. catarrhalis* as the organism. The Benjamini & Hochberg correction was applied to adjust p-values and control the False Discovery Rate (FDR) [68], setting a significance level of 0.05. The analysis considered the Biological Process (BP), Molecular Function (MF), and Cellular Component (CC) categories of Gene Ontology (GO).

### 6.5 Antibiotic resistance

Based on the CDS/sORF amino acid sequences generated by Bakta v1.8.1 [56], AMRFinderPlus v3.12.8 [69,70] was used to identify antibiotic and heavy metal resistance genes. A specific search was conducted for the *bro-1* and *bro-2* genes, which are associated with β-lactam resistance in *M. catarrhalis*. Additionally, the analysis was performed in plus mode, enabling the detection of additional genes related to resistance to biocides, metals, and other stress response mechanisms. The identification of these genes was performed using BLAST within the curated reference database of AMRFinderPlus [69,70], which includes acquired resistance elements and specific point mutations for each taxon.

### 6.6 Virulence factors

For the identification of proteins associated with virulence factors in *M. catarrhalis*, a total of 21 proteins were selected, linked to key functions in adhesion, immune system evasion, and iron acquisition. These included outer membrane proteins such as CopB (AAU43876.1 / AAU43878.1 / AAU43879.1) [71–73], LbpB (AAC31373.1) [71,73,74], M35 (AAX99225.1) [71,73,75], OmpCD (AAS75593.1) [71,73,76], OmpE (AAA64436.1) [71,73,77], OmpG1a (AAQ24464.1) [71,73,78], and OmpG1b (AAS21221.1 / AAS21227.1 / AAS21232.1) [71,73,79]; adhesion factors such as Mcap (ABM05621.1 / ABM05625.1 / ABM05628.1) [71,73,80], MclS (AGM39706.1) [71,73,81], McmA (ABL74969.1) [71,73], MhaB1 (ABQ43330.1 / ABQ43331.1) [71,73,82], MhaB2 (ABQ43328.1 / ABQ43329.1) [71,73,82], MhaC (ABQ42353.1 / ABQ42358.1) [71,73,82], MID/Hag (AAL78284.1 / AAL78285.2 / AAX56610.1) [71,73,82,83], UspA1 (AAD43469.1 / AAF36416.1 / AAN84895.1 / ACC44784.1) [71,73,84], UspA2 (AAD43468.1 / AAF40119.1 / AAN84896.1 / AAO59378.1 / AAW62383.1) [71,73,84], and UspA2H (AAF40120.1 / AAF40121.1 / AAO59379.1 / ABH07416.1) [71,73,85]; iron acquisition systems such as TbpB (AAC34274.1 / AAC34279.1) [71,73,82,86]; and proteins associated with type IV pilus formation such as PilA (AAV33390.1 / AEB33767.1) [71,73,87], PilQ (AAV33391.1) [71,73,87], and PilT (AAV33392.1) [71,73,87]. The sequences of these proteins were obtained from NCBI (https://www.ncbi.nlm.nih.gov/protein/) and used as queries in a BLASTp v2.14.0 [88] analysis against the amino acid sequences of CDS/sORFs generated by Bakta [56]. The search was performed on the proteins encoded by each assembled genome, retaining only the single best match per genome. Matches were prioritized first by e-value, followed by bitscore, percentage identity, and finally percentage coverage of the alignment [89].

### 6.7 *In silico* classification of a species

To estimate the genomic relationship among the 345 sequences, to estimate the genomic relationship among the 345 sequences, we use the Average Nucleotide Identity (ANI) between genome pairs in Pyani v0.2.12 [90] applying the ANIm method, which utilizes MUMmer for sequence alignment[91]. A threshold of ≥95% sequence identity by ANI was considered to determine that two genomes belong to the same species, as this value corresponds to the 70% identity threshold by DNA-DNA hybridization (DDH), traditionally used for the classification of prokaryotic species [90]. Additionally, *in silico* DNA-DNA hybridization (dDDH) [92–97] was applied to estimate the genomic relationship at the phylogroup level. This method is based on the same principle as experimental DNA-DNA hybridization (DDH), used for bacterial species definition. However, in this case, dDDH is *in silico,* implemented using assembled sequences. A threshold of ≥70% dDDH, along with a G+C content difference ≤1%, was established to indicate that two strains belong to the same species [98]. Using the species identified from the dDDH analysis, a phylogenomic analysis was performed following the steps described in section 3.1 Phylogenomic analysis, applying a core genome threshold of 80%. Subsequently, a new ANI calculation was conducted using Pyani v0.2.12[90], aiming to assess the genomic relationship between these species and those within the phylogroup, thereby determining their genomic proximity.

## 7. Results and Discussion

### 7.1 Pangenome composition and adaptive potential of *M. catarrhalis*

The pangenome analysis of *M. catarrhalis* revealed a total of 3,692 genes, which were classified based on their frequency in the analyzed genomes. Of these, 1,061 genes were identified as core genes, present in 100% of the genomes. These include genes involved in basic cellular and metabolic processes, growth, and development, which are essential for the survival of the species [99]. On the other hand, 2,631 genes were classified as accessory genes, subdivided into 397 soft core genes, 472 shell genes, and 1,762 cloud genes. The shell and cloud genes contribute to genetic diversity and the adaptive capacity of isolates, including environmental adaptation, drug resistance, and host adaptation [100,101]. In general, accessory genes are involved in secondary metabolism, stress response, and interactions with other organisms, and are likely associated with adaptation to specific environments [99]. The high number of accessory genes in the *M. catarrhalis* pangenome suggests significant genetic and adaptive variability in this species [102]. To assess the pangenome expansion dynamics, Heap’s Law was applied, yielding a value of γ = 0.126. This result indicates that *M. catarrhalis* has an open pangenome, which implies a high capacity for acquiring new genes, possibly through horizontal gene transfer (HGT) and the presence of various mobile genetic elements (MGEs) [103,104]. This capacity enables the species to adapt to different environments and respond to diverse selective pressures [105]. In the case of *M. catarrhalis*, the openness of its pangenome may be closely related to its ability to colonize the human respiratory tract and withstand different selective pressures, including exposure to antibiotics and the host immune response.

### 7.2 Phylogenetic clustering and lineage associations in *M. catarrhalis*

Through phylogenetic clustering, three distinct phylogroups were identified, designated as Phylogroup A, Phylogroup B, and Phylogroup C. To assess the possible correlation of these phylogroups with previously reported lineages, marker strains were used, whose classification into specific lineages had already been established in previous studies [19]. The results indicated that Phylogroup A was associated with a seroresistant lineage, whereas Phylogroup B corresponded to a serosensitive lineage, and Phylogroup C to a divergent lineage (Figure 1a). Additionally, an analysis of accessory genes was performed to evaluate their clustering structure (Figure S1). The results showed that the observed grouping matched the phylogroups defined in the core genome-based phylogeny, suggesting that both core and accessory genes reflect consistent patterns of diversification in *M. catarrhalis*.

**Figure 1.**
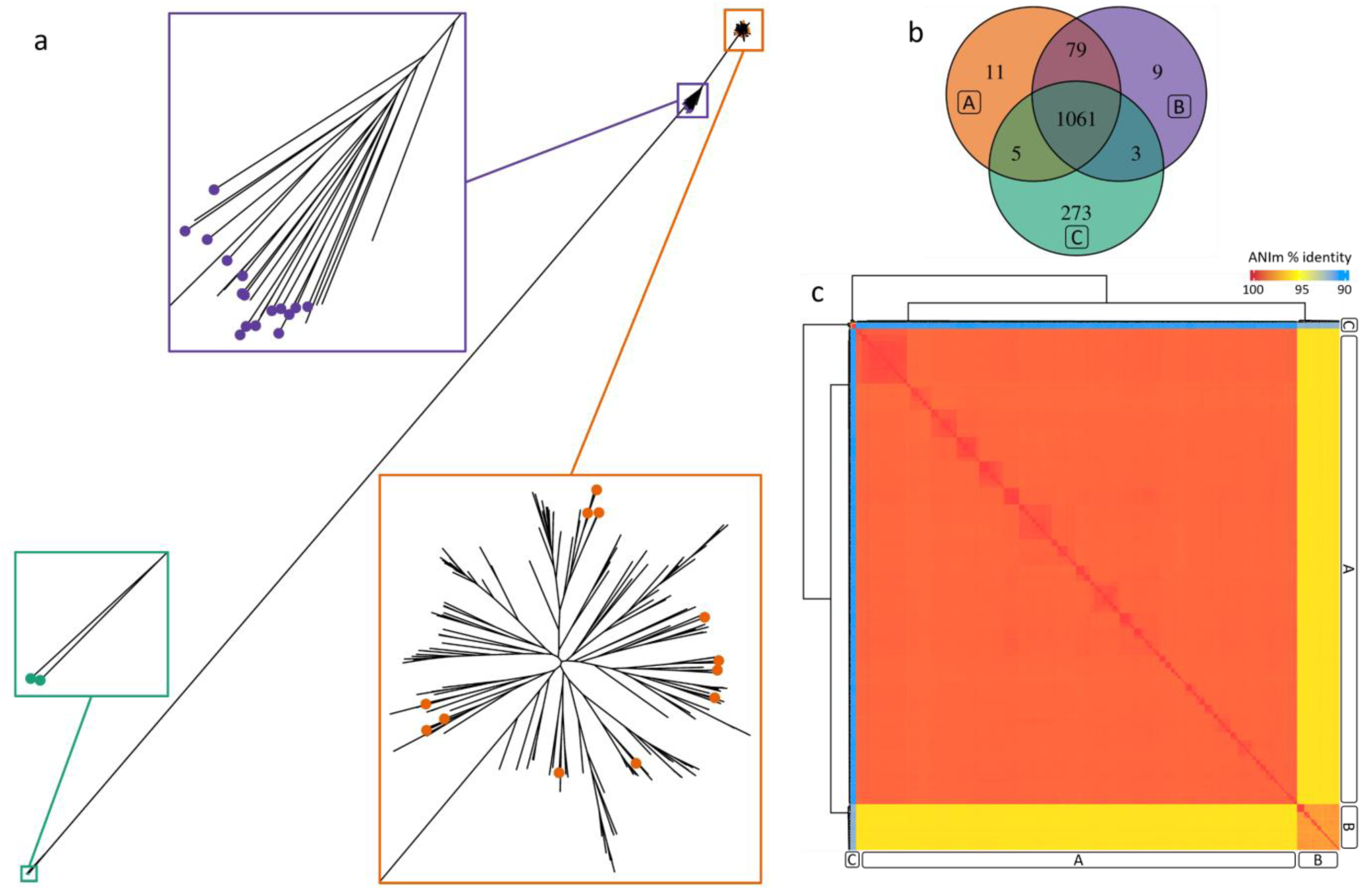
Phylogenetic and comparative genomic analysis of *M. catarrhalis*. (a) Maximum likelihood phylogenetic tree of *M. catarrhalis* strains, showing the three phylogroups: A (orange), B (purple), and C (green). Each dot on the branches represents an experimentally evaluated serotype. (b) Venn diagram displaying the number of unique and shared genes among phylogroups A, B, and C. (c) Heatmap of Average Nucleotide Identity (ANI) values calculated using Pyani, illustrating the genomic similarity among strains from different phylogroups. The letters A, B, and C correspond to the classification of phylogroups.

### 7.3 Genomic diversity and functional specialization among *M. catarrhalis* phylogroups

The analysis of the genes shared among the phylogroups (Figure 1b) of *M. catarrhalis* revealed the presence of several essential genes involved in DNA repair and replication, as well as structural integrity. Among the genes related to DNA recombination repair, *recA*, *recX*, *recC*, *recF*, *recG*, *recO*, and *recR* were identified, playing a key role in post-replicative repair [106]. RecA is essential for homologous recombination and the response to RecFOR DNA damage, facilitating the restoration of stalled replication forks due to unrepaired lesions [107]. In *Escherichia coli*, the and RecBCD pathways enable RecA loading in damaged regions [108,109], while in *Bacillus subtilis*, RecO and RecR are crucial for the formation of repair centers [110]. On the other hand, in the initiation and elongation of DNA replication, the genes *dnaA*, *dnaE*, *dnaK*, *dnaN*, and *dnaQ* were identified. DnaA is responsible for recognizing and unwinding the origin of replication (*oriC*), facilitating the loading of the DnaB helicase, in complex with DnaC, to establish the replication fork [111,112].

Regarding responses under stress, the genes *ahpC* and *ahpF* play a key role in oxidative stress resistance in *M. catarrhalis*. AhpC is a thiol-dependent peroxiredoxin that catalyzes the reduction of hydrogen peroxide (H₂O₂) [113,114], organic hydroperoxides such as tert-butyl hydroperoxide (t-BHP) and cumene hydroperoxide (CHP), as well as peroxynitrite, thereby protecting the bacteria from oxidative damage [115,116]. AhpF, on the other hand, is a flavoprotein with oxidoreductase activity that restores AhpC to its reduced form, enabling its continuous antioxidant function [114,117]. In other bacteria, such as *Bacillus subtilis*, AhpC has been identified as a general stress protein induced in response to adverse conditions, including heat, osmotic stress, and the stationary phase of growth [118].

Additionally, genes associated with the formation of type IV pili, essential structures for adhesion, motility, and genetic transformation in *M. catarrhalis*, were identified. The genes *pilA*, *pil B*, *pil C*, *pil F*, *pilO*, *pilP*, *pilQ*, *pilV* and *pilW* participate in the biogenesis and assembly of the pilus. PilA is the major subunit of the filament [87], PilQ functions as the secretin protein in the outer membrane through which the pilus is extruded [87], while PilT is the ATPase that regulates its retraction [119]. Previous studies have demonstrated that type IV pili in *M. catarrhalis* are constitutively expressed and regulated by iron via the Fur regulator, playing a crucial role in pathogenesis and host interaction. Finally, genes involved in Lipid II biosynthesis were identified, including *murA*, *murB*, *murC*, *murE*, *murF*, and *murG*, which form part of the dcw cluster (*division and cell wall*) [120]. This set of genes catalyzes the stepwise synthesis of the peptidoglycan precursor, an essential component of the bacterial cell wall [121](For more details, see Table S2).

These results suggest that *M. catarrhalis* has developed mechanisms enabling it to adapt, evade host immune responses, and maintain structural integrity, all of which are essential for successful colonization and persistence within the host. Specifically, the presence of genes associated with adhesion and biofilm formation underscores the potential of these strains to establish chronic infections, particularly in respiratory environments. Furthermore, genes related to DNA repair, replication, and stress responses indicate the bacterium’s high adaptability, facilitating survival under challenging conditions such as antibiotic exposure, oxidative stress, or host immune responses. Given these findings, proteins such as PilQ and CopB emerge as promising therapeutic targets. PilQ represents an attractive target due to its critical role in type IV pilus biogenesis and host-pathogen interactions, essential for adhesion and colonization [122]. Similarly, CopB, an outer membrane protein, serves as a significant antigenic factor involved in nutrient acquisition and immune evasion [123], making it an attractive target for vaccine development or antibody-based therapeutics.

Among the Phylogroup A and B of *M. catarrhalis*, the genes *dppB*, *dppC*, and *dppD* were identified as part of the Dpp (dipeptide permease) system [124], a transporter belonging to the ABC (ATP-binding cassette) superfamily, responsible for the uptake of dipeptides and some tripeptides into the cell [125]. In *Helicobacter pylori*, the Dpp transporter consists of five proteins: DppA, DppB, DppC, DppD, and DppF [126,127]. Within this system, DppB and DppC are membrane proteins that form the permease channel for the substrate, while DppD and DppF are cytoplasmic proteins responsible for ATP hydrolysis, driving the import of peptides into the cell [128]. Additionally, genes *moaA*, *moaB*, *moaC*, *moaD*, *moaE*, *modA*, and *modC*, associated with molybdate metabolism, were identified. Molybdate is an essential cofactor for various bacterial enzymes. In bacteria, nitrate reductases depend on molybdenum cofactors (Mo-cofactors) to catalyze the reduction of nitrate to nitrite [129], a crucial process in anaerobic respiration and nitrogen assimilation [130]. The moaABCDE operon in *E. coli* plays a fundamental role in the biosynthesis of molybdopterin (MPT), a precursor molecule required for the activation of molybdenum-dependent enzymes [131]. In this process, MoaA, MoaB, and MoaC catalyze the synthesis of precursor Z, while MPT synthase, formed by MoaD and MoaE, adds sulfur groups to form MPT [131].

These results highlight the presence of key metabolic systems exclusively in phylogroups A and B of *M. catarrhalis*, suggesting a differential adaptive capacity compared to phylogroup C. The identification of the Dpp transport system (DppB, DppC, and DppD) implies a significant metabolic advantage for peptide and nutrient acquisition [126,127], which could be associated with increased efficiency in nutrient-rich environments within the host, such as the respiratory mucosa. Likewise, the exclusive presence of the *moa* operon in these phylogroups, involved in molybdate metabolism, suggests that they may exploit molybdenum-dependent enzymes for essential processes such as anaerobic respiration or nitrogen assimilation [129]. These differences may explain variations in colonization capacity, persistence, and even virulence among phylogroups.

Additionally, the modA and modC genes encode molybdate transport proteins that facilitate the uptake of this essential metal into the cell. Between Phylogroups A and C, the genes *pstA*, *pstB*, *pstC*, and *pstS* were identified, encoding the components of the phosphate-specific transport (Pst) system [132]. PstS, a periplasmic phosphate-binding protein [133]; PstA and PstC, transmembrane proteins forming the transport channel; and PstB, a cytoplasmic protein with a nucleotide-binding domain essential for ATP hydrolysis and active phosphate transport [134]. On the other hand, between phylogroups B and C, the *ybaK* gene was identified, encoding an aminoacyl-tRNA deacylase involved in correcting errors in amino acid loading onto tRNAs [135]. In *H. influenzae*, YbaK has been characterized as a moderately specific aa-tRNA deacylase, capable of hydrolyzing mischarged Cys-tRNA^Cys^, as well as other aa-tRNAs such as Gly-tRNA^Gly^, Ala-tRNA^Ala^, Ser-tRNA^Ser^, Pro-tRNA^Pro^, and Met-tRNA^Met^ [135]. Its activity is essential for translation homeostasis and the prevention of errors in protein synthesis (For more details, see Table S2).

The different transport and metabolic systems identified among phylogroups suggest functional adaptations that may influence the pathogenic potential of *M. catarrhalis*. The exclusive presence of *modA* and *modC* in phylogroups A and B implies a greater reliance on molybdate-dependent enzymes, which could enhance survival in oxygen-limited environments, such as biofilms or inflamed tissues [136]. The Pst system in phylogroups A and C suggests alternative strategies for phosphate acquisition, a key element in bacterial virulence and stress adaptation [134]. Additionally, the presence of *ybaK* in phylogroups B and C, involved in maintaining translational fidelity, may contribute to protein homeostasis under stress conditions [135], potentially enhancing bacterial persistence. These metabolic differences could impact colonization efficiency, immune evasion, and antibiotic resistance.

The phylogroups also exhibited specific genes, in Phylogroup A, the *pgpB* gene was identified, which encodes a phosphatidylglycerophosphate phosphatase (PGPP) involved in phospholipid metabolism. In *E. coli*, the PgpB enzyme catalyzes the conversion of phosphatidylglycerophosphate to phosphatidylglycerol, a crucial step in the biosynthesis of anionic phospholipids [137]. In Phylogroup B, the *repB* gene was identified, which is involved in the regulation of extracellular enzyme and siderophore production in *Pseudomonas viridiflava* [138] The *repB* locus encodes a response regulator homologous to *gacA*, a key component of two-component regulatory systems in *Pseudomonas syringae* and *Pseudomonas fluorescens* [139]. These systems play a fundamental role in regulating the production of virulence factors, such as pectate lyases, proteases, and alginate, as well as being involved in iron acquisition through siderophores. In Phylogroup C, the *oppA*, *oppB*, *oppC*, and *oppF* genes were identified, encoding components of the oligopeptide permease (Opp) system, an ABC transporter responsible for peptide uptake [27]. In *M. catarrhalis*, this system is crucial for the acquisition of arginine, an essential amino acid for bacterial survival and adaptation [140]. Additionally, this phylogroup also contained the *argC* and *argJ* genes, which encode enzymes involved in arginine biosynthesis. In *E. coli*, the N-acetylornithine transaminase (ArgD) fulfills transamination reactions in both the arginine biosynthetic pathway and the DAP pathway for lysine [141], while in *Corynebacterium glutamicum*, two distinct enzymes (ArgD and DapC) catalyze the reactions in the two pathways, respectively [142,143] (For more details, see Table S2).

### 7.4 Phylogenetic and genomic distinctions of Phylogroup C: A potential new taxonomic entity?

The functional analysis of the genes identified in each phylogroup reveals key differences in their genomic composition. While Phylogroups A and B share a considerable number of genes and exhibit a close phylogenetic relationship, Phylogroup C is more distantly related (Figure 1b), suggesting a higher degree of evolutionary divergence [21]. The number of genes shared between Phylogroup C and the other two phylogroups is significantly lower, whereas Phylogroups A and B share a substantially greater number of genes, indicating a closer genomic relationship between them.

Furthermore, Phylogroup C exhibits a significantly higher number of unique genes compared to the other phylogroups, suggesting potential functional or adaptive differences. Additionally, the total genomic content of each phylogroup varies considerably: Phylogroup A contains 1,710±52 genes, Phylogroup B 1,737±50, and Phylogroup C 1,877±18. These differences in genetic composition, along with the low number of genes shared with the other phylogroups, suggest that Phylogroup C may represent a distinct taxonomic entity within the Moraxella genus or potentially a different species [19].

To understand the genomic relationship among the phylogroups (Figure 1c), an ANI analysis based on ANIm was performed to assess the percentage of identity between genomes. The results showed that the similarity between Phylogroups A and B was greater than 95%, indicating a high genetic relationship between them. However, the comparison of Phylogroup C with Phylogroups A and B showed values below 95%, suggesting greater genomic divergence. Based on these results, an *in silico* DNA-DNA hybridization (dDDH) (Table 1) analysis was conducted for the genomes of Phylogroup C (GCF_001656295.1, GCF_001656335.1, GCF_001656355.1, and GCF_001656375.1), which showed the highest similarity to *Moraxella canis*, with a coincidence index (C.I. d0, in %) ranging from 65.8% to 75.7%. In second place, these same genomes showed similarity to *M. catarrhalis*, with dDDH values between 53.4% and 61.0%, indicating a lower genomic relationship with this species. In both cases, a G+C content difference greater than 1% was observed, reinforcing the hypothesis that Phylogroup C exhibits significant taxonomic divergence. To compare the genomic relationship with the other phylogroups, a representative strain of Phylogroup A (GCF_000193045.1) and one from Phylogroup B (GCF_001656415.1) were included in the analysis. The results showed that the Phylogroup A strain had dDDH values between 95.0% and 97.9% with *M. catarrhalis*, confirming its classification within this species. In the case of Phylogroup B, dDDH values ranged between 81.6% and 88.4%, and the G+C content difference between Phylogroups A and B was only 0.13%, indicating a closer phylogenetic relationship between them, despite some genomic variability within *M. catarrhalis*. These results suggest that while Phylogroups A and B represent variations within *M. catarrhalis*, Phylogroup C exhibits greater genomic divergence, potentially requiring reclassification within the *Moraxella* genus [19].

**Table 1.** Digital DNA-DNA Hybridization (dDDH) values of Phylogroup C.

| Identifier | Subject strain | C.I. (d0, in %) | C.I. (d4, in %) | C.I. (d6, in %) | G+C content difference (in %) |
| --- | --- | --- | --- | --- | --- |
| GCF 001656295.1 | *M. canis* | 65.8 - 73.4 | 44.6 - 49.7 | 62.8 - 69.4 | 1.46 |
|  | *M. catarrhalis* | 53.9 - 61.0 | 34.4 - 39.4 | 49.5 - 55.7 | 2.07 |
| GCF 001656335.1 | *M. canis* | 68.1 - 75.7 | 45.0 - 50.1 | 64.8 - 71.5 | 1.39 |
|  | *M. catarrhalis* | 53.6 - 60.7 | 34.0 - 39.0 | 49.1 - 55.3 | 2.14 |
| GCF 001656355.1 | *M. canis* | 68.1 - 75.7 | 44.7 - 49.8 | 64.7 - 71.3 | 1.43 |
|  | *M. catarrhalis* | 53.4 - 60.4 | 33.8 - 38.8 | 48.9 - 55.0 | 2.09 |
| GCF 001656375.1 | *M. canis* | 68.1 - 75.7 | 45.0 - 50.2 | 64.8 - 71.5 | 1.4 |
|  | *M. catarrhalis* | 53.5 - 60.6 | 33.9 - 38.9 | 49.0 - 55.2 | 2.13 |
| GCF 000193045.1* | *M. catarrhalis* | 95.0 - 97.9 | 90.2 - 93.9 | 96.4 - 98.4 | 0.13 |
|  | *M. canis* | 37.7 - 44.5 | 24.4 - 29.2 | 33.5 - 39.5 | 3.66 |
| GCF 001656415.1* | *M. catarrhalis* | 81.6 - 88.4 | 62.4 - 68.1 | 81.2 - 87.2 | 0.13 |
|  | *M. canis* | 38.2 - 45.0 | 24.7 - 29.6 | 34.0 - 40.0 | 3.66 |

The table presents dDDH values obtained using three Genome BLAST Distance Phylogeny (GBDP) formulas: d0 (HSP length/total genome length), d4 (sum of identities in HSPs/HSP length), and d6 (sum of identities in HSPs/total genome length). The confidence intervals (C.I.) for each formula provide an estimate of variability in similarity values. \* Correspond to the representative strain of Phylogroup A (GCF\_000193045.1) and one from Phylogroup B (GCF\_001656415.1).

The phylogenetic analysis conducted with the species obtained from the dDDH results revealed that the genomes of Phylogroup C exhibit a distinct evolutionary relationship within the *Moraxella* genus. The resulting phylogenetic showed a common node from which two main branches emerge (Figure 2a), one grouping the representative strains of Phylogroups A and B, confirming their close relationship, and another that subdivides into two clades. The first clade contains exclusively the GCF_001656295.1 strain, while the second splits into two sub-branches, one containing Moraxella canis and the other comprising the genomes GCF_001656335.1, GCF_001656355.1, and GCF_001656375.1. These results indicate that while Phylogroup C shows greater phylogenetic proximity to *M. canis* than to *M. catarrhalis*, there remains significant divergence within this group, suggesting that it may represent a distinct taxonomic entity within the Moraxella genus [19]. To complement this analysis, the Average Nucleotide Identity (ANI) (Figure 2b)was calculated among the same species included in the phylogenetic analysis. The results showed that the genomes of Phylogroup C had 93% nucleotide identity with *M. canis*, while their identity with the representatives of Phylogroup A and Phylogroup B was 91% and 92%, respectively.

**Figure 2:**
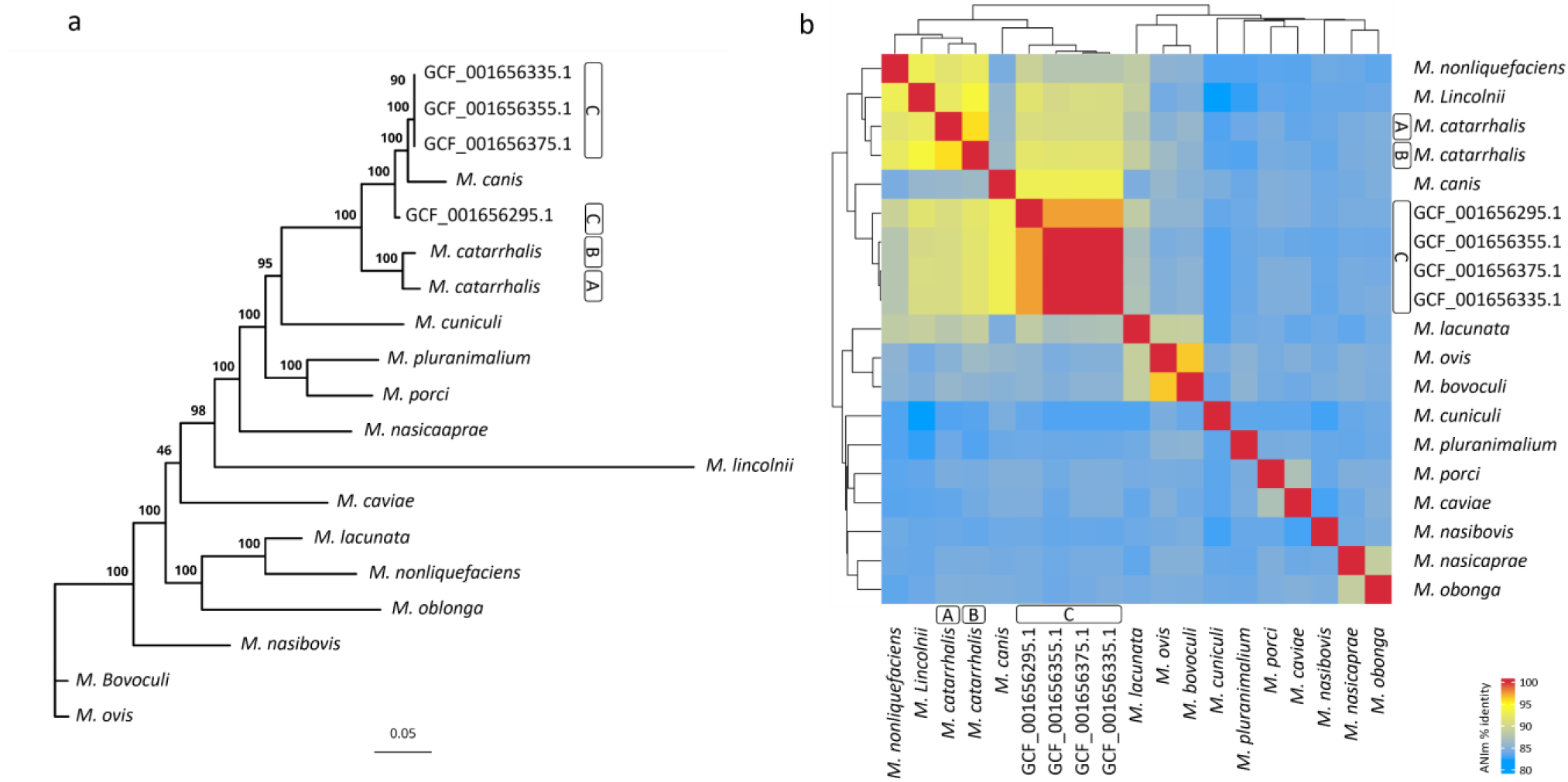
Phylogenetic and genomic differences of Phylogroup. **C.** (a) Maximum likelihood phylogenetic tree including various *Moraxella* species, highlighting the distinct positioning of Phylogroup C. (b) Heatmap of Average Nucleotide Identity (ANI) values calculated using Pyani, displaying the genomic similarity among the same *Moraxella* species analyzed in the phylogenetic tree

The distinct genomic characteristics of Phylogroup C, including its lower gene content overlap with Phylogroups A and B, its higher proportion of unique genes, and its greater genomic divergence as revealed by ANI and dDDH analyses, suggest that it may represent a separate taxonomic entity within the *Moraxella* genus. The ANI values below 95% when compared to Phylogroups A and B, along with dDDH indices closer to *M. canis* than to *M. catarrhalis*, indicate that Phylogroup C does not align with the genomic parameters of *M. catarrhalis*. Additionally, the G+C content difference exceeding 1% reinforces the hypothesis that it may belong to a different *Moraxella* species.

To determine whether Phylogroup C constitutes a novel species or a distinct lineage within an existing *Moraxella* species, further studies are required. Phenotypic characterization is essential to assess whether it exhibits distinct metabolic or structural traits. Comparative functional genomics could clarify whether its unique genes provide specific ecological adaptations. Additionally, experimental validation of gene expression and function may help define its taxonomic placement. Given its genomic divergence, Phylogroup C may represent more than just intraspecific variation and could be classified as a candidate species within *Moraxella*. However, formal classification would require meeting key taxonomic thresholds, such as dDDH values above 70% and ANI values exceeding 95% with a recognized species.

Based on the results suggesting that the members of Phylogroup C may represent a species different from *M. catarrhalis*, subsequent analyses were conducted excluding this group. Consequently, the pangenome and the corresponding phylogeny were reconstructed considering only the data from Phylogroups A and B.

### 7.5 Evaluation of genomic and epidemiological patterns

After reconstructing the pangenome, considering only Phylogroups A and B, a total of 3,219 genes were identified and classified based on their frequency in the analyzed genomes. Among these, 1,435 genes were identified as core genes, present in 100% of the genomes. Additionally, 30 soft core genes (*95% ≤ strains < 100%*), 474 shell genes (*15% ≤ strains < 95%*), and 1,280 cloud genes (*<15% of the strains*) were found. The distribution of accessory genes (shell and cloud) is detailed in Figure S2. To assess the pangenome expansion dynamics, Heap’s Law was applied, yielding a γ value of 0.093. This represents a decrease compared to the previously obtained value when Phylogroup C was included (γ = 0.126), which indicated a higher gene acquisition rate in the pangenome. The reduction in γ after removing Phylogroup C suggests that this group significantly contributed to the genetic variability of the species. These results support the hypothesis that Phylogroup C could represent a distinct species, whose inclusion increased the gene acquisition rate and, consequently, the openness of the *M. catarrhalis* pangenome.

To further understand the relationship between genomic diversity and epidemiological factors, phylogenetic analysis of Phylogroups A and B was correlated with available epidemiological data (Figure 3.), including geographic location, year of isolation, and host condition at the time of sampling. However, no clear relationship was observed between the phylogenetic structure and these variables, suggesting that the genomic diversity of *M. catarrhalis* is not strongly influenced by geographical or temporal factors. In contrast, a clear correlation was found with the previously reported seroresistant and serosensitive lineages [19,21]. Isolates belonging to Phylogroup A were predominantly associated with the seroresistant lineage, while those in Phylogroup B corresponded mainly to the serosensitive lineage. This classification aligns with previous studies that have reported differences in complement system susceptibility and virulence between these lineages.

**Figure 3:**
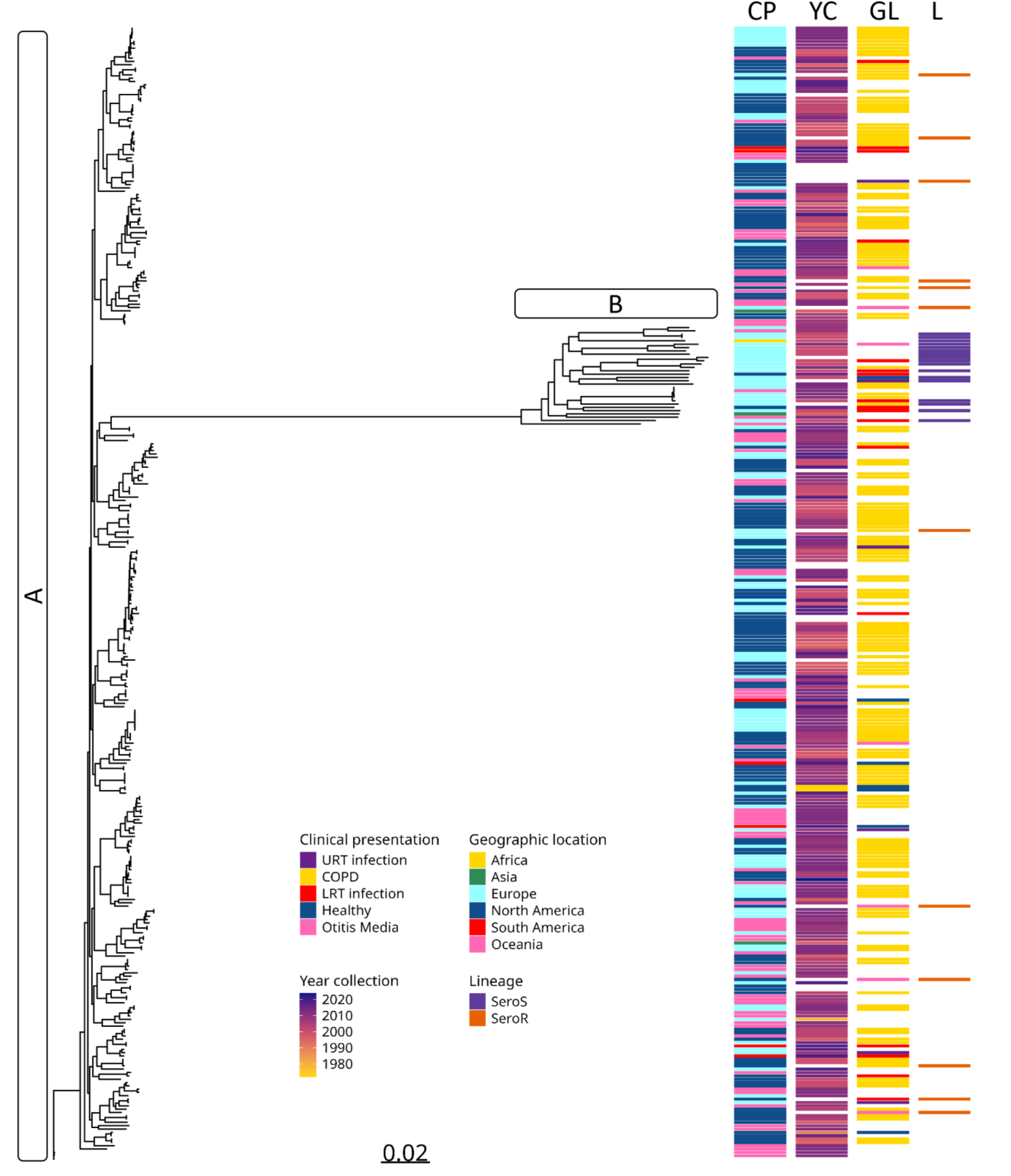
Genomic diversity and epidemiological factors of *M. catarrhalis.* Maximum-likelihood phylogenetic tree of *M. catarrhalis*, considering only Phylogroups A and B. The metadata includes clinical presentation (CP), year of collection (YC), geographic location (GL), and lineage classification (L).

### 7.6 Functional Characterization of the Core Genome of *M.catarrhalis*

The analysis of the core genes of *Moraxella catarrhalis* using Gene Ontology (GO) (Figure 4) revealed that most of these proteins are involved in essential functions for bacterial survival and adaptation, encompassing key structural, functional, and metabolic aspects.

**Figure 4:**
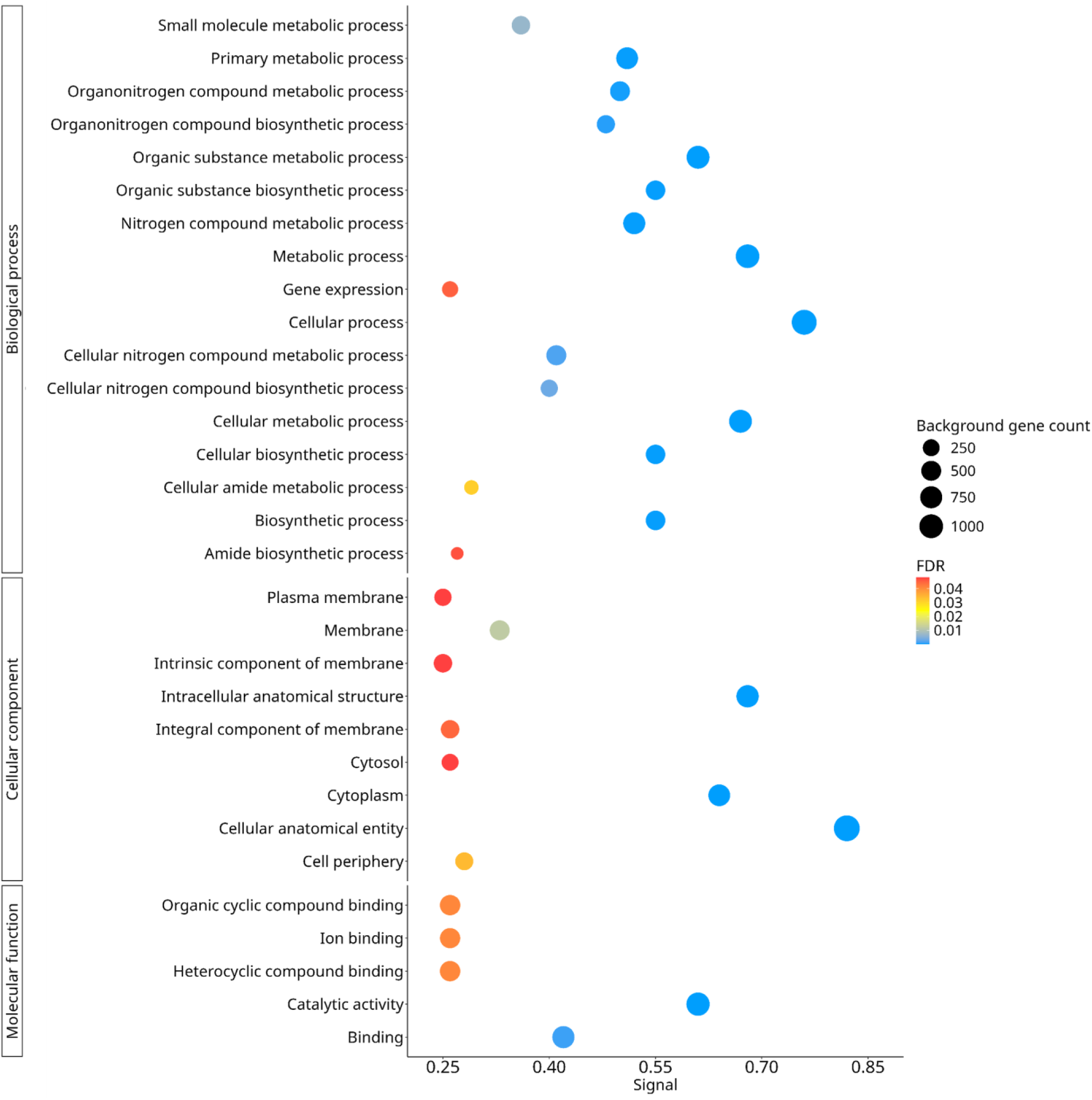
Functional analysis of the core genome of *M. catarrhalis*. Gene Ontology (GO) enrichment analysis of the core genome genes. GO terms are classified into Biological Process, Cellular Component, and Molecular Function. Each term is represented by a circle, where the size indicates the number of associated genes, and the color reflects the False Discovery Rate (FDR).

In the cellular components category, proteins related to cellular structure and organization were identified, including cellular anatomical entity (1,042 proteins) and intracellular anatomical structure (634 proteins). Likewise, proteins associated with the plasma membrane and its components were identified, including membrane (374 proteins), intrinsic component of membrane (276 proteins), integral component of membrane (274 proteins), cell periphery (253 proteins), and plasma membrane (219 proteins). These membrane proteins play key roles in epithelial surface adhesion, cell recognition, and immune system evasion [144]. Their presence in *M. catarrhalis* suggests a crucial role in host interaction and persistence in the respiratory tract. Moreover, the lateral dynamics of these membrane components regulate protein-protein and lipid-protein interactions [145–148], influencing cell signaling mechanisms and environmental adaptation. The high representation of proteins associated with the membrane highlights the importance of these processes in the biology of *M. catarrhalis*, particularly in its ability to colonize the respiratory tract and evade host immune responses [149].

In addition, proteins related to cellular structure, organization, and the membrane, were also identified, with high representation in the cytoplasm (569 proteins) and cytosol (201 proteins). Since the cytoplasm is the primary site for metabolic activity, protein synthesis, and gene regulation [150], the abundance of these proteins suggests a key role in the adaptive response and cellular viability of *M. catarrhalis*.

Some of these cytoplasmic proteins (CPs), traditionally considered intracellular, can be exported into the extracellular environment through different mechanisms, acquiring additional functions known as "moonlighting" proteins [151,152]. In *M. catarrhalis*, these proteins can be released directly or via outer membrane vesicles (OMVs), which serve as transport systems for virulence factors and play a crucial role in immune evasion [153]. OMVs contain numerous surface and periplasmic proteins, such as UspA1, UspA2, and MID (Moraxella IgD-binding protein), many of which contribute to host immune modulation by interfering with complement activation and suppressing pro-inflammatory responses [153]. For instance, UspA1 has been shown to bind carcinoembryonic antigen-related cell adhesion molecule 1 (CEACAM-1), reducing the inflammatory response triggered by Toll-like receptor 2 (TLR2) [153]. Furthermore, proteomic analyses have identified OMV-associated proteins, including CopB and OMP E, that induce immune responses, making them promising candidates for vaccine development [154]. Given their role in virulence and persistence, some of these moonlighting proteins, such as metabolic enzymes like enolase and GAPDH (glyceraldehyde-3-phosphate dehydrogenase), may act as key targets for novel therapeutic strategies against *M. catarrhalis* infections [155].

In the molecular function category, proteins related to catalytic activity (709 proteins) and molecular binding processes (572 proteins) were identified. Among these, proteins involved in heterocyclic compound binding (407 proteins), organic cyclic compound binding (407 proteins), and ion binding (394 proteins) were predominant. These binding processes play a crucial role in enzymatic activity, molecular interactions, and bacterial adaptation.

Proteins associated with catalytic activity are essential for accelerating biochemical reactions within the cell. Enzymes act as biological catalysts, increasing reaction rates by several orders of magnitude. Without enzymatic catalysis, most cellular reactions would proceed too slowly to sustain life under physiological conditions [156]. In *M. catarrhalis*, one such catalytic enzyme is the glucosyltransferase Lgt3, which plays a crucial role in lipooligosaccharide (LOS) biosynthesis [157]. LOS are glycolipid surface molecules that contribute to bacterial colonization and virulence [158]. Lgt3 contains two distinct glycosyltransferase domains (A1 and A2), each possessing a conserved DXD motif [159], which is critical for catalytic activity. The N-terminal A1 domain is responsible for adding the first β-(1-3) glucose (Glc) residue to the inner core of the LOS molecule, while the C-terminal A2 domain sequentially incorporates β-(1-4) Glc and β-(1-6) Glc, highlighting its bifunctional nature [160]. This enzymatic activity is essential for the structural integrity of LOS, which in turn influences bacterial immune evasion and host interactions.

Heterocyclic compound-binding proteins interact with molecules containing cyclic structures that include nitrogen, oxygen, or sulfur [161], which are fundamental in nucleic acid metabolism and enzymatic function. Organic cyclic compound-binding proteins participate in metabolic pathways and secondary metabolite processing, while ion-binding proteins regulate cellular homeostasis, enzymatic activity, and molecular transport across membranes [162]. The presence of these binding functions suggests that *M. catarrhalis* possesses a dynamic molecular network that enables adaptation to diverse environmental conditions and host interactions.

Among the binding proteins, the ubiquitous surface proteins UspA1 and UspA2 are relevant in *M. catarrhalis*, as they facilitate host tissue adhesion, immune evasion, and serum resistance [163]. UspA1 functions as an adhesin, interacting with carcinoembryonic antigen-related cellular adhesion molecules (CEACAMs), fibronectin, and laminin [164]. However, sequence variability among *M. catarrhalis* strains influences UspA1’s binding properties [164], with some strains exhibiting reduced or absent binding to CEACAMs and fibronectin. This variability affects host cell interactions, bacterial colonization, and immune evasion mechanisms [164]. Beyond its role in adhesion, UspA1 is implicated in complement resistance, interacting with C3 [165] and C4b [42], thereby enhancing bacterial survival within the host. Additionally, UspA1 has been associated with epithelial cell apoptosis induction and bacterial invasion of human cells [166]. The ability of UspA1 to bind multiple ligands in distinct regions suggests a complex mechanism for coordinating host interactions [164].

In the biological processes category, proteins related to cellular metabolism (685 proteins), organic compound metabolism (675 proteins), and nitrogen compound metabolism (566 proteins) were identified. Additionally, proteins associated with cellular biosynthesis (381 proteins), small molecule metabolism (297 proteins), and gene expression (176 proteins) were present.

Metabolism-associated proteins in *M. catarrhalis* include those involved in the conversion of large organic compounds, such as carbohydrates processed through catabolic pathways, into smaller products like organic acids and carbon dioxide [167]. However, unlike other bacterial pathogens, *M. catarrhalis* lacks a complete glycolytic pathway and instead relies on amino acid and organic acid metabolism as alternative energy sources [168]. Additionally, it possesses a functional glyoxylate cycle, allowing it to utilize acetate and simple carbon compounds for energy production, facilitating its adaptation to nutrient-limited environments [168].

Regarding nitrogen metabolism, *M. catarrhalis* has developed strategies that enable its persistence within the host. It expresses pathways for ammonia assimilation, including glutamate dehydrogenase and glutamate synthase-glutamine synthetase systems, although it lacks key regulatory elements such as RpoN, PII, NtrB, and NtrC, which are common in other bacteria [168,169]. Instead, *M. catarrhalis* predominantly degrades alanine, arginine, glycine, and histidine as nitrogen sources, reinforcing its metabolic flexibility and adaptive capacity to the conditions of the respiratory tract [169].

The high representation of genes involved in cellular metabolism and biosynthesis suggests that *M. catarrhalis* has developed efficient strategies to adapt to fluctuations in nutrient availability. Its dependence on alternative energy sources and the absence of carbohydrate catabolism pathways indicate an evolutionary adaptation that enhances its survival in highly competitive niches with limited resources, such as the respiratory tract.

### 7.7 β-Lactamase Mediated Resistance in *M. catarrhalis*

The antibiotic resistance analysis revealed that Phylogroup A harbored resistance genes in 51.45% of the strains, with 47.59% carrying the *bro-1* gene and 2.89% carrying *bro-2*. In Phylogroup B, the prevalence of resistance genes was significantly higher, reaching 80.00%, with 70.00% of strains harboring *bro-1* and 6.67% carrying *bro-2*.

The *bro-1* and *bro-2* genes encode BRO β-lactamases, the primary mediators of *M. catarrhalis* resistance to β-lactam antibiotics, such as amoxicillin and penicillin. While both enzymes exhibit similar substrate profiles and inhibition patterns, BRO-1 confers a higher level of resistance than BRO-2, likely due to its higher expression levels in *bro-1*-positive strains [170]. The difference in the expression of these β-lactamases appears to be associated with promoter sequence variations, where a 21-bp deletion in the *bro-2* promoter has been identified, potentially affecting its transcription [170].

Previous studies have reported that over 90% of clinical isolates of *M. catarrhalis* worldwide produce β-lactamases, with *bro-1* being more prevalent than *bro-2* [171]. Moreover, the subcellular localization of BRO β-lactamases suggests that they function as outer membrane-associated lipoproteins, which may enhance bacterial resistance to β-lactam antibiotics and contribute to persistence within the host [172].

The predominance of the *bro-1* gene in both phylogroups aligns with previous findings, highlighting its greater impact on β-lactam resistance compared to *bro-2*. Additionally, variability in *bro-1* and *bro-2* expression suggests potential differences in genetic regulation of resistance between these phylogroups, which may be influenced by selective pressures in distinct clinical environments. The association of BRO β-lactamases with the outer membrane may further contribute to resistance stability, enhancing immune evasion and bacterial persistence in the respiratory tract.

### 7.8 Identification of virulence associated proteins in *M. catarrhalis*

The identification of virulence associated proteins in *M. catarrhalis* revealed that several virulence factors are conserved across the analyzed genomes (Figure 5). Among them, M35, McaP, McmA, OmpCD, OmpG1b, PilQ, and PilT play essential roles in host adhesion, nutrient acquisition, and the formation of specialized structures for environmental interaction (For more details, see Table S3).

**Figure 5.**
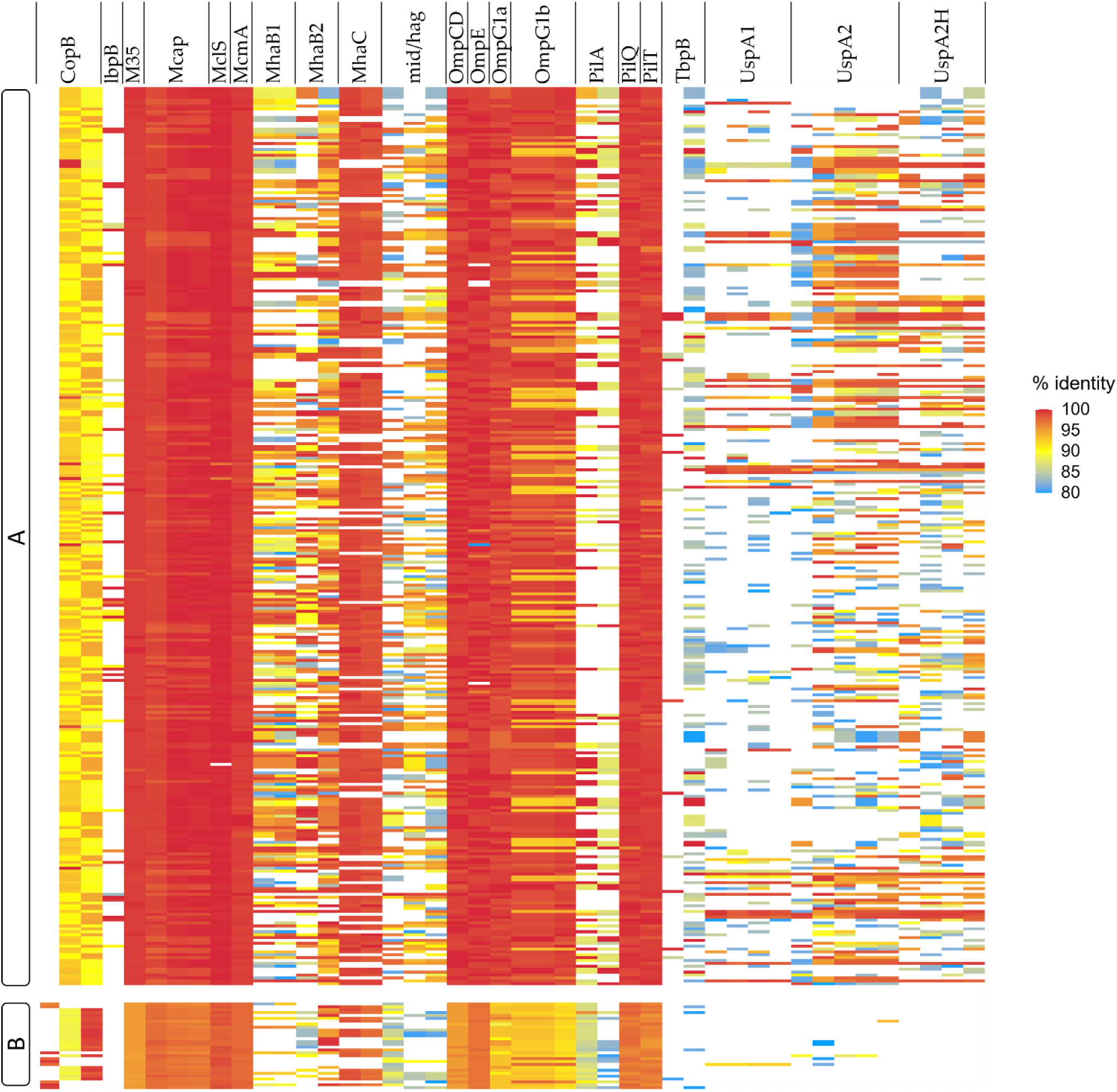
Virulence factor profiling in *M. catarrhalis*. Heatmap based on conserved virulence factors according to BLAST results, with identity and coverage percentages above 80%. Columns represent specific proteins associated with virulence, while rows correspond to different *M. catarrhalis* genomes. The letters A and B indicate the classification of the phylogroups.

Among the proteins involved in host adhesion, McaP, OmpCD, and OmpG1b were identified. McaP is an outer membrane adhesin that facilitates the attachment of *M. catarrhalis* to human epithelial cells, particularly pulmonary cells. Its N-terminal passenger domain is responsible for interacting with the cell surface, enhancing colonization stability [80]. OmpCD plays a crucial role in adhesion, as its presence significantly increases the ability of *M. catarrhalis* to bind to respiratory epithelial cells [76]. Additionally, it contributes to the structural stability of the outer membrane, which may influence bacterial resistance to adverse conditions [76]. OmpG1b, although not yet fully characterized, has been associated with membrane permeability and host interaction, suggesting a potential role in adhesion and bacterial structural stability [173].

In nutrient acquisition, the presence of outer membrane porins is essential for the uptake of molecules necessary for metabolism and survival within the host. In this process, M35 stands out as a key player. M35 is an outer membrane porin that plays a fundamental role in regulating permeability, allowing the passage of essential nutrients [75]. Its deletion negatively impacts the bacterium’s ability to survive in resource-limited environments, highlighting its importance in the metabolic adaptation of *M. catarrhalis* [75].

Finally, in the formation of specialized structures, the proteins PilQ and PilT are crucial for the biogenesis and function of type IV pili, structures essential for bacterial motility, adhesion, and genetic transformation. PilQ forms a channel in the outer membrane through which type IV pili emerge [174]. These structures enable the bacterium to adhere to cellular surfaces and form microcolonies, facilitating its persistence within the host. PilT is an ATPase responsible for the retraction of type IV pili. This process is fundamental for bacterial motility and the uptake of environmental DNA, contributing to genetic variability and bacterial adaptation to new conditions [175].

Together, these proteins play an essential role in the biology of *M. catarrhalis*, ensuring its ability to colonize the host, acquire essential nutrients, and develop specialized structures that promote its persistence and environmental adaptation. The interaction of adhesins such as McaP, OmpCD, and OmpG1b with respiratory tract epithelial cells facilitates bacterial establishment in the host, while the presence of porins like M35 enables the uptake of key metabolic molecules. Additionally, the involvement of PilQ and PilT in the biogenesis and retraction of type IV pili highlights the importance of these structures in adhesion, motility, and potential genetic transfer, contributing to the bacterium’s evolutionary plasticity. The conservation of these factors across the analyzed genomes suggests that they play critical roles in *M. catarrhalis* survival in hostile environments, underscoring their significance in pathogenesis and their ability to adapt to various ecological niches.

## 8. Conclusion

The phylogenomic analysis of *Moraxella catarrhalis* identified three phylogroups with significant genetic differences. However, ANI and dDDH results indicated that one of them exhibited considerable divergence from the other two, with values below the genomic identity thresholds established for *M. catarrhalis*. This suggests that this group could represent a new species within the *Moraxella* genus and was therefore excluded from further analyses. After its removal, the pangenome reconstruction showed a reduction in the gene acquisition rate, indicating that this divergent phylogroup was a key factor in the initially observed genetic variability.

The analysis of the two remaining phylogroups allowed for a detailed characterization of their functional differences and their association with the previously described seroresistant (Phylogroup A) and serosensitive (Phylogroup B) lineages. One of the key differences between them was the prevalence of antibiotic resistance genes. Phylogroup B exhibited a higher prevalence of resistance genes, with 80.00% of strains carrying at least one resistance determinant, including 70.00% harboring *bro-1* and 6.67% carrying *bro-2*. In contrast, Phylogroup A had a lower proportion of resistant strains, with 51.45% carrying resistance genes, including 47.59% with *bro-1* and 2.89% with *bro-2*. These findings indicate that Phylogroup B may have a greater capacity to withstand β-lactam antibiotics, while Phylogroup A might rely on other mechanisms for persistence and survival.

Both phylogroups shared key virulence factors, including outer membrane proteins such as UspA1 and UspA2, which play essential roles in immune evasion and adhesion to epithelial cells. However, they differed in transport and metabolic systems. Phylogroup A contained genes for the DppB-DppC-DppD dipeptide transport system, whereas Phylogroup B exhibited genes for the molybdate metabolism pathway, including *moaA*, *moaB*, *moaC*, *moaD*, *moaE*, *modA*, and *modC*. Additionally, AhpC, a key protein for oxidative stress resistance, was identified in both phylogroups.

These differences in genetic composition suggest that Phylogroup B may have a greater ability to resist antibiotics, while Phylogroup A exhibits adaptations related to alternative nutrient acquisition and stress response. Despite these distinctions, both phylogroups share key virulence factors, highlighting common strategies for host colonization and persistence. Given these findings, it is crucial to identify potential molecular targets that could serve as intervention points. Among them, PilQ stands out as a critical protein for type IV pilus biogenesis and host interaction, playing a fundamental role in adhesion and biofilm formation. Similarly, CopB is an important factor in iron acquisition, which is essential for bacterial survival and immune evasion. Understanding these molecular mechanisms provides valuable insights into the evolutionary dynamics of *M. catarrhalis* and its potential pathogenicity, reinforcing the need for further phylogenomic studies to explore its genetic diversity and antimicrobial resistance mechanisms.

## 9. Author statements

### 9.1 Author contributions

M.G.R, I.R-T, and J.A.U designed the project. M.G.R analysed the data, and prepared the manuscript. M.G.R, I.R-T., and J.A.U. reviewed and edited the manuscript. All authors approved the manuscript. J.A.U. supervised the study and provided funding.

## 9.4 Acknowledgements

We would like to thank the Microbial Data Science Lab at Universidad Andrés Bello for their valuable contributions to the discussion of this work.

## 10. Bibliography

1. Paykel, J.M. Moraxella (branhamella) Catarrhalis Infections. Prim. Care Update Ob. Gyns. 2002, 9, 33–35, doi:10.1016/s1068-607x(01)00099-3.

2. Spaniol, V.; Troller, R.; Schaller, A.; Aebi, C. Physiologic Cold Shock of Moraxella Catarrhalis Affects the Expression of Genes Involved in the Iron Acquisition, Serum Resistance and Immune Evasion. BMC Microbiol. 2011, 11, 182, doi:10.1186/1471-2180-11-182.

3. Del Beccaro, M.A.; Mendelman, P.M.; Inglis, A.F.; Richardson, M.A.; Duncan, N.O.; Clausen, C.R.; Stull, T.L. Bacteriology of Acute Otitis Media: A New Perspective. J. Pediatr. 1992, 120, 81–84, doi:10.1016/s0022-3476(05)80605-5.

4. Karalus, R.; Campagnari, A. Moraxella Catarrhalis: A Review of an Important Human Mucosal Pathogen. Microbes Infect. 2000, 2, 547–559, doi:10.1016/s1286-4579(00)00314-2.

5. Faden, H.; Bernstein, J.; Brodsky, L.; Stanievich, J.; Ogra, P.L. Effect of Prior Antibiotic Treatment on Middle Ear Disease in Children. Ann. Otol. Rhinol. Laryngol. 1992, 101, 87–91, doi:10.1177/000348949210100119.

6. Ruohola, A.; Meurman, O.; Nikkari, S.; Skottman, T.; Salmi, A.; Waris, M.; Osterback, R.; Eerola, E.; Allander, T.; Niesters, H.;, et al. Microbiology of Acute Otitis Media in Children with Tympanostomy Tubes: Prevalences of Bacteria and Viruses. Clin. Infect. Dis. 2006, 43, 1417–1422, doi:10.1086/509332.

7. Murphy, T.F.; Parameswaran, G.I. *Moraxella Catarrhalis*,a Human Respiratory Tract Pathogen. Clin. Infect. Dis. 2009, 49, 124–131, doi:10.1086/599375.

8. Wald, E.R. Microbiology of Acute and Chronic Sinusitis in Children and Adults. Am. J. Med. Sci. 1998, 316, 13–20, doi:10.1097/00000441-199807000-00003.

9. Brook, I.; Foote, P.A.; Hausfeld, J.N. Frequency of Recovery of Pathogens Causing Acute Maxillary Sinusitis in Adults before and after Introduction of Vaccination of Children with the 7-Valent Pneumococcal Vaccine. J. Med. Microbiol. 2006, 55, 943–946, doi:10.1099/jmm.0.46346-0.

10. Murphy, T.F.; Brauer, A.L.; Grant, B.J.B.; Sethi, S. Moraxella Catarrhalis in Chronic Obstructive Pulmonary Disease: Burden of Disease and Immune Response. Am. J. Respir. Crit. Care Med. 2005, 172, 195–199, doi:10.1164/rccm.200412-1747OC.

11. Doern, G.V. Branhamella Catarrhalis--an Emerging Human Pathogen. Diagn. Microbiol. Infect. Dis. 1986, 4, 191–201, doi:10.1016/0732-8893(86)90098-2.

12. Catlin, B.W. Branhamella Catarrhalis: An Organism Gaining Respect as a Pathogen. Clin. Microbiol. Rev. 1990, 3, 293–320, doi:10.1128/CMR.3.4.293.

13. Daoud, A.; Abuekteish, F.; Masaadeh, H. Neonatal Meningitis due to Moraxella Catarrhalis and Review of the Literature. Ann. Trop. Paediatr. 1996, 16, 199–201, doi:10.1080/02724936.1996.11747826.

14. Verduin, C.M.; Hol, C.; Fleer, A.; van Dijk, H.; van Belkum, A. Moraxella Catarrhalis: From Emerging to Established Pathogen. Clin. Microbiol. Rev. 2002, 15, 125–144, doi:10.1128/CMR.15.1.125-144.2002.

15. Patterson, T.F.; Patterson, J.E.; Masecar, B.L.; Barden, G.E.; Hierholzer, W.J., Jr; Zervos, M.J. A Nosocomial Outbreak of Branhamella Catarrhalis Confirmed by Restriction Endonuclease Analysis. J. Infect. Dis. 1988, 157, 996–1001, doi:10.1093/infdis/157.5.996.

16. Richards, S.J.; Greening, A.P.; Enright, M.C.; Morgan, M.G.; McKenzie, H. Outbreak of Moraxella Catarrhalis in a Respiratory Unit. Thorax 1993, 48, 91–92, doi:10.1136/thx.48.1.91.

17. Bootsma, H.J.; van der Heide, H.G.; van de Pas, S.; Schouls, L.M.; Mooi, F.R. Analysis of Moraxella Catarrhalis by DNA Typing: Evidence for a Distinct Subpopulation Associated with Virulence Traits. J. Infect. Dis. 2000, 181, 1376–1387, doi:10.1086/315374.

18. Pingault, N.M.; Lehmann, D.; Bowman, J.; Riley, T.V. A Comparison of Molecular Typing Methods for Moraxella Catarrhalis. J. Appl. Microbiol. 2007, 103, 2489–2495, doi:10.1111/j.1365-2672.2007.03536.x.

19. Earl, J.P.; de Vries, S.P.W.; Ahmed, A.; Powell, E.; Schultz, M.P.; Hermans, P.W.M.; Hill, D.J.; Zhou, Z.; Constantinidou, C.I.; Hu, F.Z.;, et al. Comparative Genomic Analyses of the Moraxella Catarrhalis Serosensitive and Seroresistant Lineages Demonstrate Their Independent Evolution. Genome Biol. Evol. 2016, 8, 955–974, doi:10.1093/gbe/evw039.

20. Meier, P.S.; Troller, R.; Heiniger, N.; Grivea, I.N.; Syrogiannopoulos, G.A.; Aebi, C. Moraxella Catarrhalis Strains with Reduced Expression of the UspA Outer Membrane Proteins Belong to a Distinct Subpopulation. Vaccine 2005, 23, 2000–2008, doi:10.1016/j.vaccine.2004.09.036.

21. Wirth, T.; Morelli, G.; Kusecek, B.; van Belkum, A.; van der Schee, C.; Meyer, A.; Achtman, M. The Rise and Spread of a New Pathogen: Seroresistant Moraxella Catarrhalis. Genome Res. 2007, 17, 1647–1656, doi:10.1101/gr.6122607.

22. Verhaegh, S.J.C.; Streefland, A.; Dewnarain, J.K.; Farrell, D.J.; van Belkum, A.; Hays, J.P. Age-Related Genotypic and Phenotypic Differences in Moraxella Catarrhalis Isolates from Children and Adults Presenting with Respiratory Disease in 2001-2002. Microbiology 2008, 154, 1178–1184, doi:10.1099/mic.0.2007/015057-0.

23. Shi, W.; Wen, D.; Chen, C.; Yuan, L.; Gao, W.; Tang, P.; Cheng, X.; Yao, K. β-Lactamase Production and Antibiotic Susceptibility Pattern of Moraxella Catarrhalis Isolates Collected from Two County Hospitals in China. BMC Microbiol. 2018, 18, doi:10.1186/s12866-018-1217-5.

24. Raveendran, S.; Kumar, G.; Sivanandan, R.N.; Dias, M. Moraxella Catarrhalis: A Cause of Concern with Emerging Resistance and Presence of BRO Beta-Lactamase Gene-Report from a Tertiary Care Hospital in South India. Int. J. Microbiol. 2020, 2020, 7316257, doi:10.1155/2020/7316257.

25. Deshpande, L.M.; Sader, H.S.; Fritsche, T.R.; Jones, R.N. Contemporary Prevalence of BRO Beta-Lactamases in Moraxella Catarrhalis: Report from the SENTRY Antimicrobial Surveillance Program (North America, 1997 to 2004). J. Clin. Microbiol. 2006, 44, 3775–3777, doi:10.1128/JCM.00456-06.

26. Khan, M.A.; Northwood, J.B.; Levy, F.; Verhaegh, S.J.C.; Farrell, D.J.; Van Belkum, A.; Hays, J.P. Bro {beta}-Lactamase and Antibiotic Resistances in a Global Cross-Sectional Study of Moraxella Catarrhalis from Children and Adults. J. Antimicrob. Chemother. 2010, 65, 91–97, doi:10.1093/jac/dkp401.

27. de Vries, S.P.W.; van Hijum, S.A.F.T.; Schueler, W.; Riesbeck, K.; Hays, J.P.; Hermans, P.W.M.; Bootsma, H.J. Genome Analysis of Moraxella Catarrhalis Strain BBH18, [corrected] a Human Respiratory Tract Pathogen. J. Bacteriol. 2010, 192, 3574–3583, doi:10.1128/JB.00121-10.

28. Spaniol, V.; Bernhard, S.; Aebi, C. Moraxella Catarrhalis AcrAB-OprM Efflux Pump Contributes to Antimicrobial Resistance and Is Enhanced during Cold Shock Response. Antimicrob. Agents Chemother. 2015, 59, 1886–1894, doi:10.1128/AAC.03727-14.

29. Jetter, M.; Spaniol, V.; Troller, R.; Aebi, C. Down-Regulation of Porin M35 in Moraxella Catarrhalis by Aminopenicillins and Environmental Factors and Its Potential Contribution to the Mechanism of Resistance to Aminopenicillins. J. Antimicrob. Chemother. 2010, 65, 2089–2096, doi:10.1093/jac/dkq312.

30. Hall-Stoodley, L.; Hu, F.Z.; Gieseke, A.; Nistico, L.; Nguyen, D.; Hayes, J.; Forbes, M.; Greenberg, D.P.; Dice, B.; Burrows, A.;, et al. Direct Detection of Bacterial Biofilms on the Middle-Ear Mucosa of Children with Chronic Otitis Media. JAMA 2006, 296, 202–211, doi:10.1001/jama.296.2.202.

31. Matejka, K.M.; Bremer, P.J.; Tompkins, G.R.; Brooks, H.J.L. Antibiotic Susceptibility of Moraxella Catarrhalis Biofilms in a Continuous Flow Model. Diagn. Microbiol. Infect. Dis. 2012, 74, 394–398, doi:10.1016/j.diagmicrobio.2012.08.021.

32. Budhani, R.K.; Struthers, J.K. Interaction of Streptococcus Pneumoniae and Moraxella Catarrhalis: Investigation of the Indirect Pathogenic Role of Beta-Lactamase-Producing Moraxellae by Use of a Continuous-Culture Biofilm System. Antimicrob. Agents Chemother. 1998, 42, 2521–2526, doi:10.1128/AAC.42.10.2521.

33. Høiby, N.; Bjarnsholt, T.; Givskov, M.; Molin, S.; Ciofu, O. Antibiotic Resistance of Bacterial Biofilms. Int. J. Antimicrob. Agents 2010, 35, 322–332, doi:10.1016/j.ijantimicag.2009.12.011.

34. Su, Y.-C.; Singh, B.; Riesbeck, K. Moraxella Catarrhalis: From Interactions with the Host Immune System to Vaccine Development. Future Microbiol. 2012, 7, 1073–1100, doi:10.2217/fmb.12.80.

35. Su, Y.-C.; Hallström, B.M.; Bernhard, S.; Singh, B.; Riesbeck, K. Impact of Sequence Diversity in the Moraxella Catarrhalis UspA2/UspA2H Head Domain on Vitronectin Binding and Antigenic Variation. Microbes Infect. 2013, 15, 375–387, doi:10.1016/j.micinf.2013.02.004.

36. Hoiczyk, E.; Roggenkamp, A.; Reichenbecher, M.; Lupas, A.; Heesemann, J. Structure and Sequence Analysis of Yersinia YadA and Moraxella UspAs Reveal a Novel Class of Adhesins. EMBO J. 2000, 19, 5989–5999, doi:10.1093/emboj/19.22.5989.

37. Hallström, T.; Nordström, T.; Tan, T.T.; Manolov, T.; Lambris, J.D.; Isenman, D.E.; Zipfel, P.F.; Blom, A.M.; Riesbeck, K. Immune Evasion of Moraxella Catarrhalis Involves Ubiquitous Surface Protein A-Dependent C3d Binding. J. Immunol. 2011, 186, 3120–3129, doi:10.4049/jimmunol.1002621.

38. Meier, P.S.; Troller, R.; Grivea, I.N.; Syrogiannopoulos, G.A.; Aebi, C. The Outer Membrane Proteins UspA1 and UspA2 of Moraxella Catarrhalis Are Highly Conserved in Nasopharyngeal Isolates from Young Children. Vaccine 2002, 20, 1754–1760, doi:10.1016/s0264-410x(02)00030-0.

39. Lafontaine, E.R.; Cope, L.D.; Aebi, C.; Latimer, J.L.; McCracken, G.H., Jr; Hansen, E.J. The UspA1 Protein and a Second Type of UspA2 Protein Mediate Adherence of Moraxella Catarrhalis to Human Epithelial Cells in Vitro. J. Bacteriol. 2000, 182, 1364–1373, doi:10.1128/JB.182.5.1364-1373.2000.

40. Attia, A.S.; Lafontaine, E.R.; Latimer, J.L.; Aebi, C.; Syrogiannopoulos, G.A.; Hansen, E.J. The UspA2 Protein of Moraxella Catarrhalis Is Directly Involved in the Expression of Serum Resistance. Infect. Immun. 2005, 73, 2400–2410, doi:10.1128/IAI.73.4.2400-2410.2005.

41. Pearson, M.M.; Hansen, E.J. Identification of Gene Products Involved in Biofilm Production by Moraxella Catarrhalis ETSU-9 in Vitro. Infect. Immun. 2007, 75, 4316–4325, doi:10.1128/IAI.01347-06.

42. Nordström, T.; Blom, A.M.; Forsgren, A.; Riesbeck, K. The Emerging Pathogen Moraxella Catarrhalis Interacts with Complement Inhibitor C4b Binding Protein through Ubiquitous Surface Proteins A1 and A2. J. Immunol. 2004, 173, 4598–4606, doi:10.4049/jimmunol.173.7.4598.

43. Attia, A.S.; Ram, S.; Rice, P.A.; Hansen, E.J. Binding of Vitronectin by the Moraxella Catarrhalis UspA2 Protein Interferes with Late Stages of the Complement Cascade. Infect. Immun. 2006, 74, 1597–1611, doi:10.1128/IAI.74.3.1597-1611.2006.

44. Genome. Bethesda (MD): National Library of Medicine (US), National Center for Biotechnology. Available online: https://www.ncbi.nlm.nih.gov/datasets/genome/ (accessed on 30 March 2024).

45. SRA. Bethesda (MD): National Library of Medicine (US), National Center for Biotechnology. Available online: https://www.ncbi.nlm.nih.gov/sra/ (accessed on 30 March 2024).

46. Chen, S.; Zhou, Y.; Chen, Y.; Gu, J. Fastp: An Ultra-Fast All-in-One FASTQ Preprocessor. Bioinformatics 2018, 34, i884–i890, doi:10.1093/bioinformatics/bty560.

47. Chen, S. Ultrafast One-Pass FASTQ Data Preprocessing, Quality Control, and Deduplication Using Fastp. Imeta 2023, 2, e107, doi:10.1002/imt2.107.

48. Wood, D.E.; Lu, J.; Langmead, B. Improved Metagenomic Analysis with Kraken 2. bioRxiv 2019.

49. Lu, J.; Rincon, N.; Wood, D.E.; Breitwieser, F.P.; Pockrandt, C.; Langmead, B.; Salzberg, S.L.; Steinegger, M. Metagenome Analysis Using the Kraken Software Suite. Nat. Protoc. 2022, 17, 2815– 2839, doi:10.1038/s41596-022-00738-y.

50. Prjibelski, A.; Antipov, D.; Meleshko, D.; Lapidus, A.; Korobeynikov, A. Using SPAdes DE Novo Assembler. Curr. Protoc. Bioinformatics 2020, 70, e102, doi:10.1002/cpbi.102.

51. Chklovski, A.; Parks, D.H.; Woodcroft, B.J.; Tyson, G.W. CheckM2: A Rapid, Scalable and Accurate Tool for Assessing Microbial Genome Quality Using Machine Learning. Nat. Methods 2023, 20, 1203–1212, doi:10.1038/s41592-023-01940-w.

52. Jolley, K.A.; Bliss, C.M.; Bennett, J.S.; Bratcher, H.B.; Brehony, C.; Colles, F.M.; Wimalarathna, H.; Harrison, O.B.; Sheppard, S.K.; Cody, A.J.;, et al. Ribosomal Multilocus Sequence Typing: Universal Characterization of Bacteria from Domain to Strain. Microbiology 2012, 158, 1005–1015, doi:10.1099/mic.0.055459-0.

53. Jolley, K.A.; Bray, J.E.; Maiden, M.C.J. Open-Access Bacterial Population Genomics: BIGSdb Software, the PubMLST.org Website and Their Applications. Wellcome Open Res. 2018, 3, 124, doi:10.12688/wellcomeopenres.14826.1.

54. Seemann, T. Mlst: :id: Scan Contig Files against PubMLST Typing Schemes; Github;.

55. Davie, J.J.; Earl, J.; de Vries, S.P.W.; Ahmed, A.; Hu, F.Z.; Bootsma, H.J.; Stol, K.; Hermans, P.W.M.; Wadowsky, R.M.; Ehrlich, G.D.;, et al. Comparative Analysis and Supragenome Modeling of Twelve Moraxella Catarrhalis Clinical Isolates. BMC Genomics 2011, 12, 70, doi:10.1186/1471-2164-12-70.

56. Schwengers, O.; Jelonek, L.; Dieckmann, M.A.; Beyvers, S.; Blom, J.; Goesmann, A. Bakta: Rapid and Standardized Annotation of Bacterial Genomes via Alignment-Free Sequence Identification. Microb. Genom. 2021, 7, doi:10.1099/mgen.0.000685.

57. Tonkin-Hill, G.; MacAlasdair, N.; Ruis, C.; Weimann, A.; Horesh, G.; Lees, J.A.; Gladstone, R.A.; Lo, S.; Beaudoin, C.; Floto, R.A.;, et al. Producing Polished Prokaryotic Pangenomes with the Panaroo Pipeline. Genome Biol. 2020, 21, 180, doi:10.1186/s13059-020-02090-4.

58. Stamatakis, A. RAxML Version 8: A Tool for Phylogenetic Analysis and Post-Analysis of Large Phylogenies. Bioinformatics 2014, 30, 1312–1313, doi:10.1093/bioinformatics/btu033.

59. Flouri, T.; Izquierdo-Carrasco, F.; Darriba, D.; Aberer, A.J.; Nguyen, L.-T.; Minh, B.Q.; Von Haeseler, A.; Stamatakis, A. The Phylogenetic Likelihood Library. Syst. Biol. 2015, 64, 356–362, doi:10.1093/sysbio/syu084.

60. Kozlov, A.M.; Aberer, A.J.; Stamatakis, A. ExaML Version 3: A Tool for Phylogenomic Analyses on Supercomputers. Bioinformatics 2015, 31, 2577–2579, doi:10.1093/bioinformatics/btv184.

61. Scholl, C.; Kobert, K.; Flouri, T.; Stamatakis, A. The Divisible Load Balance Problem with Shared Cost and Its Application to Phylogenetic Inference. In Proceedings of the 2016 IEEE International Parallel and Distributed Processing Symposium Workshops (IPDPSW); IEEE, May 2016.

62. Balaban, M.; Moshiri, N.; Mai, U.; Jia, X.; Mirarab, S. TreeCluster: Clustering Biological Sequences Using Phylogenetic Trees. PLoS One 2019, 14, e0221068, doi:10.1371/journal.pone.0221068.

63. Alam, S.; Dobbie, G.; Koh, Y.S.; Riddle, P.; Ur Rehman, S. Research on Particle Swarm Optimization Based Clustering: A Systematic Review of Literature and Techniques. Swarm Evol. Comput. 2014, 17, 1–13, doi:10.1016/j.swevo.2014.02.001.

64. Zhang, B.; Ren, H.; Wang, X.; Han, C.; Jin, Y.; Hu, X.; Shi, R.; Li, C.; Wang, Y.; Li, Y.;, et al. Comparative Genomics Analysis to Explore the Biodiversity and Mining Novel Target Genes of Listeria Monocytogenes Strains from Different Regions. Front. Microbiol. 2024, 15, 1424868, doi:10.3389/fmicb.2024.1424868.

65. Tettelin, H.; Riley, D.; Cattuto, C.; Medini, D. Comparative Genomics: The Bacterial Pan-Genome. Curr. Opin. Microbiol. 2008, 11, 472–477, doi:10.1016/j.mib.2008.09.006.

66. Heap_Law_for_Roary: This python3 Program Calculates the Heap Law for Roary Output; Github. Available online: https://github.com/SethCommichaux/Heap_Law_for_Roary (accessed on 27 April 2024).

67. Szklarczyk, D.; Kirsch, R.; Koutrouli, M.; Nastou, K.; Mehryary, F.; Hachilif, R.; Gable, A.L.; Fang, T.; Doncheva, N.T.; Pyysalo, S.;, et al. The STRING Database in 2023: Protein-Protein Association Networks and Functional Enrichment Analyses for Any Sequenced Genome of Interest. Nucleic Acids Res. 2023, 51, D638–D646, doi:10.1093/nar/gkac1000.

68. Benjamini, Y.; Hochberg, Y. Controlling the False Discovery Rate: A Practical and Powerful Approach to Multiple Testing. J. R. Stat. Soc. Series B Stat. Methodol. 1995, 57, 289–300, doi:10.1111/j.2517-6161.1995.tb02031.x.

69. Feldgarden, M.; Brover, V.; Haft, D.H.; Prasad, A.B.; Slotta, D.J.; Tolstoy, I.; Tyson, G.H.; Zhao, S.; Hsu, C.-H.; McDermott, P.F.;, et al. Validating the AMRFinder Tool and Resistance Gene Database by Using Antimicrobial Resistance Genotype-Phenotype Correlations in a Collection of Isolates. Antimicrob. Agents Chemother. 2019, 63, doi:10.1128/AAC.00483-19.

70. Feldgarden, M.; Brover, V.; Gonzalez-Escalona, N.; Frye, J.G.; Haendiges, J.; Haft, D.H.; Hoffmann, M.; Pettengill, J.B.; Prasad, A.B.; Tillman, G.E.;, et al. AMRFinderPlus and the Reference Gene Catalog Facilitate Examination of the Genomic Links among Antimicrobial Resistance, Stress Response, and Virulence. Sci. Rep. 2021, 11, 12728, doi:10.1038/s41598-021-91456-0.

71. Blakeway, L.V.; Tan, A.; Peak, I.R.A.; Seib, K.L. Virulence Determinants of Moraxella Catarrhalis: Distribution and Considerations for Vaccine Development. Microbiology 2017, 163, 1371–1384, doi:10.1099/mic.0.000523.

72. Chan, C.; Ng, D.; Schryvers, A.B. The Role of the Moraxella Catarrhalis CopB Protein in Facilitating Iron Acquisition from Human Transferrin and Lactoferrin. Front. Microbiol. 2021, 12, 714815, doi:10.3389/fmicb.2021.714815.

73. Morris, D.E.; Osman, K.L.; Cleary, D.W.; Clarke, S.C. The Characterization of Moraxella Catarrhalis Carried in the General Population. Microb. Genom. 2022, 8, doi:10.1099/mgen.0.000820.

74. Bonnah, R.A.; Wong, H.; Loosmore, S.M.; Schryvers, A.B. Characterization of Moraxella (Branhamella) Catarrhalis lbpB, lbpA, and Lactoferrin Receptor orf3 Isogenic Mutants. Infect. Immun. 1999, 67, 1517–1520, doi:10.1128/IAI.67.3.1517-1520.1999.

75. Easton, D.M.; Maier, E.; Benz, R.; Foxwell, A.R.; Cripps, A.W.; Kyd, J.M. Moraxella Catarrhalis M35 Is a General Porin That Is Important for Growth under Nutrient-Limiting Conditions and in the Nasopharynges of Mice. J. Bacteriol. 2008, 190, 7994–8002, doi:10.1128/JB.01039-08.

76. Holm, M.M.; Vanlerberg, S.L.; Foley, I.M.; Sledjeski, D.D.; Lafontaine, E.R. The Moraxella Catarrhalis Porin-like Outer Membrane Protein CD Is an Adhesin for Human Lung Cells. Infect. Immun. 2004, 72, 1906–1913, doi:10.1128/IAI.72.4.1906-1913.2004.

77. Murphy, T.F.; Brauer, A.L.; Yuskiw, N.; Hiltke, T.J. Antigenic Structure of Outer Membrane Protein E of Moraxella Catarrhalis and Construction and Characterization of Mutants. Infect. Immun. 2000, 68, 6250–6256, doi:10.1128/IAI.68.11.6250-6256.2000.

78. Adlowitz, D.G.; Sethi, S.; Cullen, P.; Adler, B.; Murphy, T.F. Human Antibody Response to Outer Membrane Protein G1a, a Lipoprotein of Moraxella Catarrhalis. Infect. Immun. 2005, 73, 6601– 6607, doi:10.1128/IAI.73.10.6601-6607.2005.

79. Ren, D.; Pichichero, M.E. Vaccine Targets against Moraxella Catarrhalis. Expert Opin. Ther. Targets 2016, 20, 19–33, doi:10.1517/14728222.2015.1081686.

80. Lipski, S.L.; Akimana, C.; Timpe, J.M.; Wooten, R.M.; Lafontaine, E.R. The Moraxella Catarrhalis Autotransporter McaP Is a Conserved Surface Protein That Mediates Adherence to Human Epithelial Cells through Its N-Terminal Passenger Domain. Infect. Immun. 2007, 75, 314–324, doi:10.1128/IAI.01330-06.

81. Buskirk, S.W.; Lafontaine, E.R. Moraxella Catarrhalis Expresses a Cardiolipin Synthase That Impacts Adherence to Human Epithelial Cells. J. Bacteriol. 2014, 196, 107–120, doi:10.1128/JB.00298-13.

82. de Vries, S.P.W.; Bootsma, H.J.; Hays, J.P.; Hermans, P.W.M. Molecular Aspects of Moraxella Catarrhalis Pathogenesis. Microbiol. Mol. Biol. Rev. 2009, 73, 389–406, Table of Contents, doi:10.1128/MMBR.00007-09.

83. Holm, M.M.; Vanlerberg, S.L.; Sledjeski, D.D.; Lafontaine, E.R. The Hag Protein of Moraxella Catarrhalis Strain O35E Is Associated with Adherence to Human Lung and Middle Ear Cells. Infect. Immun. 2003, 71, 4977–4984, doi:10.1128/IAI.71.9.4977-4984.2003.

84. Tan, T.T.; Nordström, T.; Forsgren, A.; Riesbeck, K. The Respiratory Pathogen Moraxella Catarrhalis Adheres to Epithelial Cells by Interacting with Fibronectin through Ubiquitous Surface Proteins A1 and A2. J. Infect. Dis. 2005, 192, 1029–1038, doi:10.1086/432759.

85. Wang, W.; Pearson, M.M.; Attia, A.S.; Blick, R.J.; Hansen, E.J. A UspA2H-Negative Variant of Moraxella Catarrhalis Strain O46E Has a Deletion in a Homopolymeric Nucleotide Repeat Common to uspA2H Genes. Infect. Immun. 2007, 75, 2035–2045, doi:10.1128/IAI.00609-06.

86. Myers, L.E.; Yang, Y.P.; Du, R.P.; Wang, Q.; Harkness, R.E.; Schryvers, A.B.; Klein, M.H.; Loosmore, S.M. The Transferrin Binding Protein B of Moraxella Catarrhalis Elicits Bactericidal Antibodies and Is a Potential Vaccine Antigen. Infect. Immun. 1998, 66, 4183–4192, doi:10.1128/IAI.66.9.4183-4192.1998.

87. Luke, N.R.; Howlett, A.J.; Shao, J.; Campagnari, A.A. Expression of Type IV Pili by Moraxella Catarrhalis Is Essential for Natural Competence and Is Affected by Iron Limitation. Infect. Immun. 2004, 72, 6262–6270, doi:10.1128/IAI.72.11.6262-6270.2004.

88. Camacho, C.; Boratyn, G.M.; Joukov, V.; Vera Alvarez, R.; Madden, T.L. ElasticBLAST: Accelerating Sequence Search via Cloud Computing. BMC Bioinformatics 2023, 24, 117, doi:10.1186/s12859-023-05245-9.

89. Fassler, J.; Cooper, P. *BLAST Glossary*; National Center for Biotechnology Information (US), 2011;.

90. Pritchard, L.; Glover, R.H.; Humphris, S.; Elphinstone, J.G.; Toth, I.K. Genomics and Taxonomy in Diagnostics for Food Security: Soft-Rotting Enterobacterial Plant Pathogens. Anal. Methods 2016, 8, 12–24, doi:10.1039/c5ay02550h.

91. Kurtz, S.; Phillippy, A.; Delcher, A.L.; Smoot, M.; Shumway, M.; Antonescu, C.; Salzberg, S.L. Versatile and Open Software for Comparing Large Genomes. Genome Biol. 2004, 5, R12, doi:10.1186/gb-2004-5-2-r12.

92. Meier-Kolthoff, J.P.; Auch, A.F.; Klenk, H.-P.; Göker, M. Genome Sequence-Based Species Delimitation with Confidence Intervals and Improved Distance Functions. BMC Bioinformatics 2013, 14, 60, doi:10.1186/1471-2105-14-60.

93. Meier-Kolthoff, J.P.; Hahnke, R.L.; Petersen, J.; Scheuner, C.; Michael, V.; Fiebig, A.; Rohde, C.; Rohde, M.; Fartmann, B.; Goodwin, L.A.;, et al. Complete Genome Sequence of DSM 30083(T), the Type Strain (U5/41(T)) of Escherichia Coli, and a Proposal for Delineating Subspecies in Microbial Taxonomy. Stand. Genomic Sci. 2014, 9, 2, doi:10.1186/1944-3277-9-2.

94. Lefort, V.; Desper, R.; Gascuel, O. FastME 2.0: A Comprehensive, Accurate, and Fast Distance-Based Phylogeny Inference Program. Mol. Biol. Evol. 2015, 32, 2798–2800, doi:10.1093/molbev/msv150.

95. Kreft, Ł.; Botzki, A.; Coppens, F.; Vandepoele, K.; Van Bel, M. PhyD3: A Phylogenetic Tree Viewer with Extended phyloXML Support for Functional Genomics Data Visualization. Bioinformatics 2017, 33, 2946–2947, doi:10.1093/bioinformatics/btx324.

96. Meier-Kolthoff, J.P.; Göker, M. TYGS Is an Automated High-Throughput Platform for State-of-the-Art Genome-Based Taxonomy. Nat. Commun. 2019, 10, 2182, doi:10.1038/s41467-019-10210-3.

97. Meier-Kolthoff, J.P.; Carbasse, J.S.; Peinado-Olarte, R.L.; Göker, M. TYGS and LPSN: A Database Tandem for Fast and Reliable Genome-Based Classification and Nomenclature of Prokaryotes. Nucleic Acids Res.2022, 50, D801–D807, doi:10.1093/nar/gkab902.

98. Meier-Kolthoff, J.P.; Klenk, H.-P.; Göker, M. Taxonomic Use of DNA G+C Content and DNA-DNA Hybridization in the Genomic Age. Int. J. Syst. Evol. Microbiol. 2014, 64, 352–356, doi:10.1099/ijs.0.056994-0.

99. Švara, A.; Sun, H.; Fei, Z.; Khan, A. Advancing Apple Genetics Research: Malus Coronaria and Malus Ioensis Genomes and a Gene Family-Based Pangenome of Native North American Apples. DNA Res. 2024, 31, doi:10.1093/dnares/dsae026.

100. Medini, D.; Donati, C.; Tettelin, H.; Masignani, V.; Rappuoli, R. The Microbial Pan-Genome. Curr. Opin. Genet. Dev. 2005, 15, 589–594, doi:10.1016/j.gde.2005.09.006.

101. Kumari, K.; Rawat, V.; Shadan, A.; Sharma, P.K.; Deb, S.; Singh, R.P. In-Depth Genome and Pan-Genome Analysis of a Metal-Resistant Bacterium Pseudomonas Parafulva OS-1. Front. Microbiol. 2023, 14, 1140249, doi:10.3389/fmicb.2023.1140249.

102. Segerman, B. The Genetic Integrity of Bacterial Species: The Core Genome and the Accessory Genome, Two Different Stories. Front. Cell. Infect. Microbiol. 2012, 2, 116, doi:10.3389/fcimb.2012.00116.

103. Bosi, E.; Fondi, M.; Orlandini, V.; Perrin, E.; Maida, I.; de Pascale, D.; Tutino, M.L.; Parrilli, E.; Lo Giudice, A.; Filloux, A.;, et al. The Pangenome of (Antarctic) Pseudoalteromonas Bacteria: Evolutionary and Functional Insights. BMC Genomics 2017, 18, 93, doi:10.1186/s12864-016-3382-y.

104. Yin, Z.; Liang, J.; Zhang, M.; Chen, B.; Yu, Z.; Tian, X.; Deng, X.; Peng, L. Pan-Genome Insights into Adaptive Evolution of Bacterial Symbionts in Mixed Host-Microbe Symbioses Represented by Human Gut Microbiota Bacteroides Cellulosilyticus. Sci. Total Environ. 2024, 927, 172251, doi:10.1016/j.scitotenv.2024.172251.

105. Gaba, S.; Kumari, A.; Medema, M.; Kaushik, R. Pan-Genome Analysis and Ancestral State Reconstruction of Class Halobacteria: Probability of a New Super-Order. Sci. Rep. 2020, 10, 21205, doi:10.1038/s41598-020-77723-6.

106. Webb, B.L.; Cox, M.M.; Inman, R.B. Recombinational DNA Repair: The RecF and RecR Proteins Limit the Extension of RecA Filaments beyond Single-Strand DNA Gaps. Cell 1997, 91, 347–356, doi:10.1016/s0092-8674(00)80418-3.

107. Lenhart, J.S.; Brandes, E.R.; Schroeder, J.W.; Sorenson, R.J.; Showalter, H.D.; Simmons, L.A. RecO and RecR Are Necessary for RecA Loading in Response to DNA Damage and Replication Fork Stress. J. Bacteriol. 2014, 196, 2851–2860, doi:10.1128/JB.01494-14.

108. Kowalczykowski, S.C.; Dixon, D.A.; Eggleston, A.K.; Lauder, S.D.; Rehrauer, W.M. Biochemistry of Homologous Recombination in Escherichia Coli. Microbiol. Rev. 1994, 58, 401–465, doi:10.1128/mr.58.3.401-465.1994.

109. Cox, M.M. Regulation of Bacterial RecA Protein Function. Crit. Rev. Biochem. Mol. Biol. 2007, 42, 41–63, doi:10.1080/10409230701260258.

110. Lenhart, J.S.; Schroeder, J.W.; Walsh, B.W.; Simmons, L.A. DNA Repair and Genome Maintenance in Bacillus Subtilis. Microbiol. Mol. Biol. Rev. 2012, 76, 530–564, doi:10.1128/MMBR.05020-11.

111. Chodavarapu, S.; Kaguni, J.M. Replication Initiation in Bacteria. Enzymes 2016, 39, 1–30, doi:10.1016/bs.enz.2016.03.001.

112. Kaguni, J. The Macromolecular Machines That Duplicate the Escherichia Coli Chromosome as Targets for Drug Discovery. Antibiotics (Basel) 2018, 7, 23, doi:10.3390/antibiotics7010023.

113. den Hengst, C.D.; Buttner, M.J. Redox Control in Actinobacteria. Biochim. Biophys. Acta 2008, 1780, 1201–1216, doi:10.1016/j.bbagen.2008.01.008.

114. Si, M.; Hu, M.; Yang, M.; Peng, Z.; Li, D.; Zhao, Y. Characterization of Oxidative Stress-Induced Cgahp, a Gene Coding for Alkyl Hydroperoxide Reductase, from Industrial Importance Corynebacterium Glutamicum. Research Square 2023.

115. Bryk, R.; Griffin, P.; Nathan, C. Peroxynitrite Reductase Activity of Bacterial Peroxiredoxins. Nature 2000, 407, 211–215, doi:10.1038/35025109.

116. Zhang, B.; Gu, H.; Yang, Y.; Bai, H.; Zhao, C.; Si, M.; Su, T.; Shen, X. Molecular Mechanisms of AhpC in Resistance to Oxidative Stress in Burkholderia Thailandensis. Front. Microbiol. 2019, 10, 1483, doi:10.3389/fmicb.2019.01483.

117. Poole, L.B. Bacterial Defenses against Oxidants: Mechanistic Features of Cysteine-Based Peroxidases and Their Flavoprotein Reductases. Arch. Biochem. Biophys. 2005, 433, 240–254, doi:10.1016/j.abb.2004.09.006.

118. Antelmann, H.; Engelmann, S.; Schmid, R.; Hecker, M. General and Oxidative Stress Responses in Bacillus Subtilis: Cloning, Expression, and Mutation of the Alkyl Hydroperoxide Reductase Operon. J. Bacteriol. 1996, 178, 6571–6578, doi:10.1128/jb.178.22.6571-6578.1996.

119. Adams, D.W.; Pereira, J.M.; Stoudmann, C.; Stutzmann, S.; Blokesch, M. The Type IV Pilus Protein PilU Functions as a PilT-Dependent Retraction ATPase. PLoS Genet. 2019, 15, e1008393, doi:10.1371/journal.pgen.1008393.

120. Real, G.; Henriques, A.O. Localization of the Bacillus Subtilis murB Gene within the Dcw Cluster Is Important for Growth and Sporulation. J. Bacteriol. 2006, 188, 1721–1732, doi:10.1128/JB.188.5.1721-1732.2006.

121. Smith, C.A. Structure, Function and Dynamics in the Mur Family of Bacterial Cell Wall Ligases. J. Mol. Biol. 2006, 362, 640–655, doi:10.1016/j.jmb.2006.07.066.

122. Leighton, T.L.; Buensuceso, R.N.C.; Howell, P.L.; Burrows, L.L. Biogenesis of Pseudomonas Aeruginosa Type IV Pili and Regulation of Their Function: Pseudomonas Aeruginosatype IV Pili. Environ. Microbiol. 2015, 17, 4148–4163, doi:10.1111/1462-2920.12849.

123. Helminen, M.E.; Beach, R.; Maciver, I.; Jarosik, G.; Hansen, E.J.; Leinonen, M. Human Immune Response against Outer Membrane Proteins of Moraxella (Branhamella) Catarrhalis Determined by Immunoblotting and Enzyme Immunoassay. Clin. Diagn. Lab. Immunol. 1995, 2, 35–39, doi:10.1128/cdli.2.1.35-39.1995.

124. Garai, P.; Chandra, K.; Chakravortty, D. Bacterial Peptide Transporters: Messengers of Nutrition to Virulence. Virulence 2017, 8, 297–309, doi:10.1080/21505594.2016.1221025.

125. Paulsen, I.T.; Skurray, R.A. The POT Family of Transport Proteins. Trends Biochem. Sci. 1994, 19, 404, doi:10.1016/0968-0004(94)90087-6.

126. Davis, G.S.; Mobley, H.L.T. Contribution of dppA to Urease Activity in Helicobacter Pylori 26695. Helicobacter 2005, 10, 416–423, doi:10.1111/j.1523-5378.2005.00348.x.

127. Weinberg, M.V.; Maier, R.J. Peptide Transport in Helicobacter Pylori: Roles of Dpp and Opp Systems and Evidence for Additional Peptide Transporters. J. Bacteriol. 2007, 189, 3392–3402, doi:10.1128/JB.01636-06.

128. Xu, X.; Chen, J.; Huang, X.; Feng, S.; Zhang, X.; She, F.; Wen, Y. The Role of a Dipeptide Transporter in the Virulence of Human Pathogen, Helicobacter Pylori. Front. Microbiol. 2021, 12, 633166, doi:10.3389/fmicb.2021.633166.

129. Rubio, L.M.; Flores, E.; Herrero, A. Molybdopterin Guanine Dinucleotide Cofactor in Synechococcus Sp. Nitrate Reductase: Identification of mobA and Isolation of a Putative moeB Gene. FEBS Lett. 1999, 462, 358–362, doi:10.1016/s0014-5793(99)01556-2.

130. Wootton, J.C.; Nicolson, R.E.; Cock, J.M.; Walters, D.E.; Burke, J.F.; Doyle, W.A.; Bray, R.C. Enzymes Depending on the Pterin Molybdenum Cofactor: Sequence Families, Spectroscopic Properties of Molybdenum and Possible Cofactor-Binding Domains. Biochim. Biophys. Acta 1991, 1057, 157–185, doi:10.1016/s0005-2728(05)80100-8.

131. Rajagopalan, K.V. In: Escherichia Coli and Salmonella: Cellular and Molecular Biology. In Escherichia coli and Salmonella: Cellular and Molecular Biology ; Neidhardt, F.C., Ed, F.C., Eds.; 1996; pp. 674–679.

132. Amemura, M.; Makino, K.; Shinagawa, H.; Kobayashi, A.; Nakata, A. Nucleotide Sequence of the Genes Involved in Phosphate Transport and Regulation of the Phosphate Regulon in Escherichia Coli. J. Mol. Biol. 1985, 184, 241–250, doi:10.1016/0022-2836(85)90377-8.

133. Gerdes, R.G.; Rosenberg, H. The Relationship between the Phosphate-Binding Protein and a Regulator Gene Product from Escherichia Coli. Biochim. Biophys. Acta 1974, 351, 77–86, doi:10.1016/0005-2795(74)90066-x.

134. Higgins, C.F.; Hiles, I.D.; Whalley, K.; Jamieson, D.J. Nucleotide Binding by Membrane Components of Bacterial Periplasmic Binding Protein-Dependent Transport Systems. EMBO J. 1985, 4, 1033– 1039, doi:10.1002/j.1460-2075.1985.tb03735.x.

135. Ruan, B.; Söll, D. The Bacterial YbaK Protein Is a Cys-tRNAPro and Cys-tRNA Cys Deacylase. J. Biol. Chem. 2005, 280, 25887–25891, doi:10.1074/jbc.M502174200.

136. Pederick, V.G.; Eijkelkamp, B.A.; Ween, M.P.; Begg, S.L.; Paton, J.C.; McDevitt, C.A. Acquisition and Role of Molybdate in Pseudomonas Aeruginosa. Appl. Environ. Microbiol. 2014, 80, 6843–6852, doi:10.1128/aem.02465-14.

137. Funk, C.R.; Zimniak, L.; Dowhan, W. The pgpA and pgpB Genes of Escherichia Coli Are Not Essential: Evidence for a Third Phosphatidylglycerophosphate Phosphatase. J. Bacteriol. 1992, 174, 205–213, doi:10.1128/jb.174.1.205-213.1992.

138. Poole, K.; McKay, G.A. Iron Acquisition and Its Control in Pseudomonas Aeruginosa: Many Roads Lead to Rome. Front. Biosci. 2003, 8, d661–d686, doi:10.2741/1051.

139. Liao, C.H.; McCallus, D.E.; Wells, J.M.; Tzean, S.S.; Kang, G.Y. The repB Gene Required for Production of Extracellular Enzymes and Fluorescent Siderophores in Pseudomonas Viridiflava Is an Analog of the gacA Gene of Pseudomonas Syringae. Can. J. Microbiol. 1996, 42, 177–182, doi:10.1139/m96-026.

140. Jones, M.M.; Johnson, A.; Koszelak-Rosenblum, M.; Kirkham, C.; Brauer, A.L.; Malkowski, M.G.; Murphy, T.F. Role of the Oligopeptide Permease ABC Transporter of Moraxella Catarrhalis in Nutrient Acquisition and Persistence in the Respiratory Tract. Infect. Immun. 2014, 82, 4758–4766, doi:10.1128/IAI.02185-14.

141. Ledwidge, R.; Blanchard, J.S. The Dual Biosynthetic Capability of N-Acetylornithine Aminotransferase in Arginine and Lysine Biosynthesis. Biochemistry 1999, 38, 3019–3024, doi:10.1021/bi982574a.

142. Hartmann, M.; Tauch, A.; Eggeling, L.; Bathe, B.; Möckel, B.; Pühler, A.; Kalinowski, J. Identification and Characterization of the Last Two Unknown Genes, dapC and dapF, in the Succinylase Branch of the L-Lysine Biosynthesis of Corynebacterium Glutamicum. J. Biotechnol. 2003, 104, 199–211, doi:10.1016/s0168-1656(03)00156-1.

143. Wu, Y.; Zhang, J.; Wang, B.; Zhang, Y.; Li, H.; Liu, Y.; Yin, J.; He, D.; Luo, H.; Gan, F.;, et al. Dissecting the Arginine and Lysine Biosynthetic Pathways and Their Relationship in Haloarchaeon Natrinema Gari J7-2 via Endogenous CRISPR-Cas System-Based Genome Editing. Microbiol. Spectr. 2023, 11, e0028823, doi:10.1128/spectrum.00288-23.

144. Foster, T.J.; Geoghegan, J.A.; Ganesh, V.K.; Höök, M. Adhesion, Invasion and Evasion: The Many Functions of the Surface Proteins of Staphylococcus Aureus. Nat. Rev. Microbiol. 2014, 12, 49–62, doi:10.1038/nrmicro3161.

145. Zwaal, R.F.; Comfurius, P.; Bevers, E.M. Lipid-Protein Interactions in Blood Coagulation. Biochim. Biophys. Acta 1998, 1376, 433–453, doi:10.1016/s0304-4157(98)00018-5.

146. Laganowsky, A.; Reading, E.; Allison, T.M.; Ulmschneider, M.B.; Degiacomi, M.T.; Baldwin, A.J.; Robinson, C.V. Membrane Proteins Bind Lipids Selectively to Modulate Their Structure and Function. Nature 2014, 510, 172–175, doi:10.1038/nature13419.

147. Whited, A.M.; Johs, A. The Interactions of Peripheral Membrane Proteins with Biological Membranes. Chem. Phys. Lipids 2015, 192, 51–59, doi:10.1016/j.chemphyslip.2015.07.015.

148. Muller, M.P.; Jiang, T.; Sun, C.; Lihan, M.; Pant, S.; Mahinthichaichan, P.; Trifan, A.; Tajkhorshid, E. Characterization of Lipid-Protein Interactions and Lipid-Mediated Modulation of Membrane Protein Function through Molecular Simulation. Chem. Rev. 2019, 119, 6086–6161, doi:10.1021/acs.chemrev.8b00608.

149. Siegel, S.J.; Weiser, J.N. Mechanisms of Bacterial Colonization of the Respiratory Tract. Annu. Rev. Microbiol. 2015, 69, 425–444, doi:10.1146/annurev-micro-091014-104209.

150. Parry, B.R.; Surovtsev, I.V.; Cabeen, M.T.; O’Hern, C.S.; Dufresne, E.R.; Jacobs-Wagner, C. The Bacterial Cytoplasm Has Glass-like Properties and Is Fluidized by Metabolic Activity. Cell 2014, 156, 183–194, doi:10.1016/j.cell.2013.11.028.

151. Ebner, P.; Prax, M.; Nega, M.; Koch, I.; Dube, L.; Yu, W.; Rinker, J.; Popella, P.; Flötenmeyer, M.; Götz, F. Excretion of Cytoplasmic Proteins (ECP) in Staphylococcus Aureus. Mol. Microbiol. 2015, 97, 775–789, doi:10.1111/mmi.13065.

152. Ebner, P.; Götz, F. Bacterial Excretion of Cytoplasmic Proteins (ECP): Occurrence, Mechanism, and Function. Trends Microbiol. 2019, 27, 176–187, doi:10.1016/j.tim.2018.10.006.

153. Schaar, V.; de Vries, S.P.W.; Perez Vidakovics, M.L.A.; Bootsma, H.J.; Larsson, L.; Hermans, P.W.M.; Bjartell, A.; Mörgelin, M.; Riesbeck, K. Multicomponent Moraxella Catarrhalis Outer Membrane Vesicles Induce an Inflammatory Response and Are Internalized by Human Epithelial Cells. Cell. Microbiol. 2011, 13, 432–449, doi:10.1111/j.1462-5822.2010.01546.x.

154. Augustyniak, D.; Seredyński, R.; McClean, S.; Roszkowiak, J.; Roszniowski, B.; Smith, D.L.; Drulis-Kawa, Z.; Mackiewicz, P. Virulence Factors of Moraxella Catarrhalis Outer Membrane Vesicles Are Major Targets for Cross-Reactive Antibodies and Have Adapted during Evolution. Sci. Rep. 2018, 8, 4955, doi:10.1038/s41598-018-23029-7.

155. Karner, A.; Gesslbauer, B.; Spreitzer, A.; Almer, J.; Smidt, M.; Schüler, W.; Fartmann, B.; Zimmermann, W.; Meinke, A.; Kungl, A.J. Profiling the Membrane and Glycosaminoglycan-Binding Proteomes of *Moraxella Catarrhalis*. J. Proteome Res. 2016, 15, 3055–3097, doi:10.1021/acs.jproteome.6b00187.

156. Cooper, G.M. The Central Role of Enzymes as Biological Catalysts; Sinauer Associates, 2000;.

157. Edwards, K.J.; Allen, S.; Gibson, B.W.; Campagnari, A.A. Characterization of a Cluster of Three Glycosyltransferase Enzymes Essential for Moraxella Catarrhalis Lipooligosaccharide Assembly. J. Bacteriol. 2005, 187, 2939–2947, doi:10.1128/JB.187.9.2939-2947.2005.

158. Erridge, C.; Bennett-Guerrero, E.; Poxton, I.R. Structure and Function of Lipopolysaccharides. Microbes Infect. 2002, 4, 837–851, doi:10.1016/s1286-4579(02)01604-0.

159. Jing, W.; DeAngelis, P.L. Analysis of the Two Active Sites of the Hyaluronan Synthase and the Chondroitin Synthase of Pasteurella Multocida. Glycobiology 2003, 13, 661–671, doi:10.1093/glycob/cwg085.

160. Luke-Marshall, N.R.; Edwards, K.J.; Sauberan, S.; St Michael, F.; Vinogradov, E.V.; Cox, A.D.; Campagnari, A.A. Characterization of a Trifunctional Glucosyltransferase Essential for Moraxella Catarrhalis Lipooligosaccharide Assembly. Glycobiology 2013, 23, 1013–1021, doi:10.1093/glycob/cwt042.

161. Borah, P.; Hazarika, S.; Chettri, A.; Sharma, D.; Deka, S.; Venugopala, K.N.; Shinu, P.; Al-Shar’i, N.A.; Bardaweel, S.K.; Deb, P.K. Heterocyclic Compounds as Antimicrobial Agents. In Viral, Parasitic, Bacterial, and Fungal Infections; Elsevier, 2023; pp. 781–804 ISBN 9780323857307.

162. Assmann, S.M.; Armstrong, F. Hormonal Regulation of Ion Transporters: The Guard Cell System. In Biochemistry and Molecular Biology of Plant Hormones; New comprehensive biochemistry; Elsevier, 1999; pp. 337–361 ISBN 9780444898258.

163. Spaniol, V.; Heiniger, N.; Troller, R.; Aebi, C. Outer Membrane Protein UspA1 and Lipooligosaccharide Are Involved in Invasion of Human Epithelial Cells by Moraxella Catarrhalis. Microbes Infect. 2008, 10, 3–11, doi:10.1016/j.micinf.2007.09.014.

164. Brooks, M.J.; Sedillo, J.L.; Wagner, N.; Wang, W.; Attia, A.S.; Wong, H.; Laurence, C.A.; Hansen, E.J.; Gray-Owen, S.D. Moraxella Catarrhalis Binding to Host Cellular Receptors Is Mediated by Sequence-Specific Determinants Not Conserved among All UspA1 Protein Variants. Infect. Immun. 2008, 76, 5322–5329, doi:10.1128/IAI.00572-08.

165. Nordström, T.; Blom, A.M.; Tan, T.T.; Forsgren, A.; Riesbeck, K. Ionic Binding of C3 to the Human Pathogen Moraxella Catarrhalis Is a Unique Mechanism for Combating Innate Immunity. J. Immunol. 2005, 175, 3628–3636, doi:10.4049/jimmunol.175.6.3628.

166. N’Guessan, P.D.; Vigelahn, M.; Bachmann, S.; Zabel, S.; Opitz, B.; Schmeck, B.; Hippenstiel, S.; Zweigner, J.; Riesbeck, K.; Singer, B.B.;, et al. The UspA1 Protein of*Moraxella Catarrhalis* induces CEACAM-1–dependent Apoptosis in Alveolar Epithelial Cells. J. Infect. Dis. 2007, 195, 1651–1660, doi:10.1086/514820.

167. Muñoz-Elías, E.J.; McKinney, J.D. Carbon Metabolism of Intracellular Bacteria. Cell. Microbiol. 2006, 8, 10–22, doi:10.1111/j.1462-5822.2005.00648.x.

168. Wang, W.; Reitzer, L.; Rasko, D.A.; Pearson, M.M.; Blick, R.J.; Laurence, C.; Hansen, E.J. Metabolic Analysis of Moraxella Catarrhalis and the Effect of Selected in Vitro Growth Conditions on Global Gene Expression. Infect. Immun. 2007, 75, 4959–4971, doi:10.1128/IAI.00073-07.

169. Juni, E.; Heym, G.A.; Avery, M. Defined Medium for Moraxella (Branhamella) Catarrhalis. Appl. Environ. Microbiol. 1986, 52, 546–551, doi:10.1128/aem.52.3.546-551.1986.

170. Schmitz, F.-J.; Beeck, A.; Perdikouli, M.; Boos, M.; Mayer, S.; Scheuring, S.; Köhrer, K.; Verhoef, J.; Fluit, A.C. Production of BRO β-Lactamases and Resistance to Complement in European *Moraxella Catarrhalis* Isolates. J. Clin. Microbiol. 2002, 40, 1546–1548, doi:10.1128/jcm.40.4.1546-1548.2002.

171. Esel, D.; Ay-Altintop, Y.; Yagmur, G.; Gokahmetoglu, S.; Sumerkan, B. Evaluation of Susceptibility Patterns and BRO Beta-Lactamase Types among Clinical Isolates of Moraxella Catarrhalis. Clin. Microbiol. Infect. 2007, 13, 1023–1025, doi:10.1111/j.1469-0691.2007.01776.x.

172. Bootsma, H.J.; Aerts, P.C.; Posthuma, G.; Harmsen, T.; Verhoef, J.; van Dijk, H.; Mooi, F.R. Moraxella (Branhamella)catarrhalis BRO β-Lactamase: A Lipoprotein of Gram-Positive Origin? J. Bacteriol. 1999, 181, 5090–5093, doi:10.1128/jb.181.16.5090-5093.1999.

173. Adlowitz, D.G.; Kirkham, C.; Sethi, S.; Murphy, T.F. Human Serum and Mucosal Antibody Responses to Outer Membrane Protein G1b of Moraxella Catarrhalis in Chronic Obstructive Pulmonary Disease. FEMS Immunol. Med. Microbiol. 2006, 46, 139–146, doi:10.1111/j.1574-695X.2005.00020.x.

174. Assalkhou, R.; Balasingham, S.; Collins, R.F.; Frye, S.A.; Davidsen, T.; Benam, A.V.; Bjørås, M.; Derrick, J.P.; Tønjum, T. The Outer Membrane Secretin PilQ from Neisseria Meningitidis Binds DNA. Microbiology 2007, 153, 1593–1603, doi:10.1099/mic.0.2006/004200-0.

175. Forest, K.T.; Satyshur, K.A.; Worzalla, G.A.; Hansen, J.K.; Herdendorf, T.J. The Pilus-Retraction Protein PilT: Ultrastructure of the Biological Assembly. Acta Crystallogr. D Biol. Crystallogr. 2004, 60, 978–982, doi:10.1107/S0907444904006055.

