## Supplementary material for "A global perspective on the genomics of Moraxella catarrhalis": Figure S1

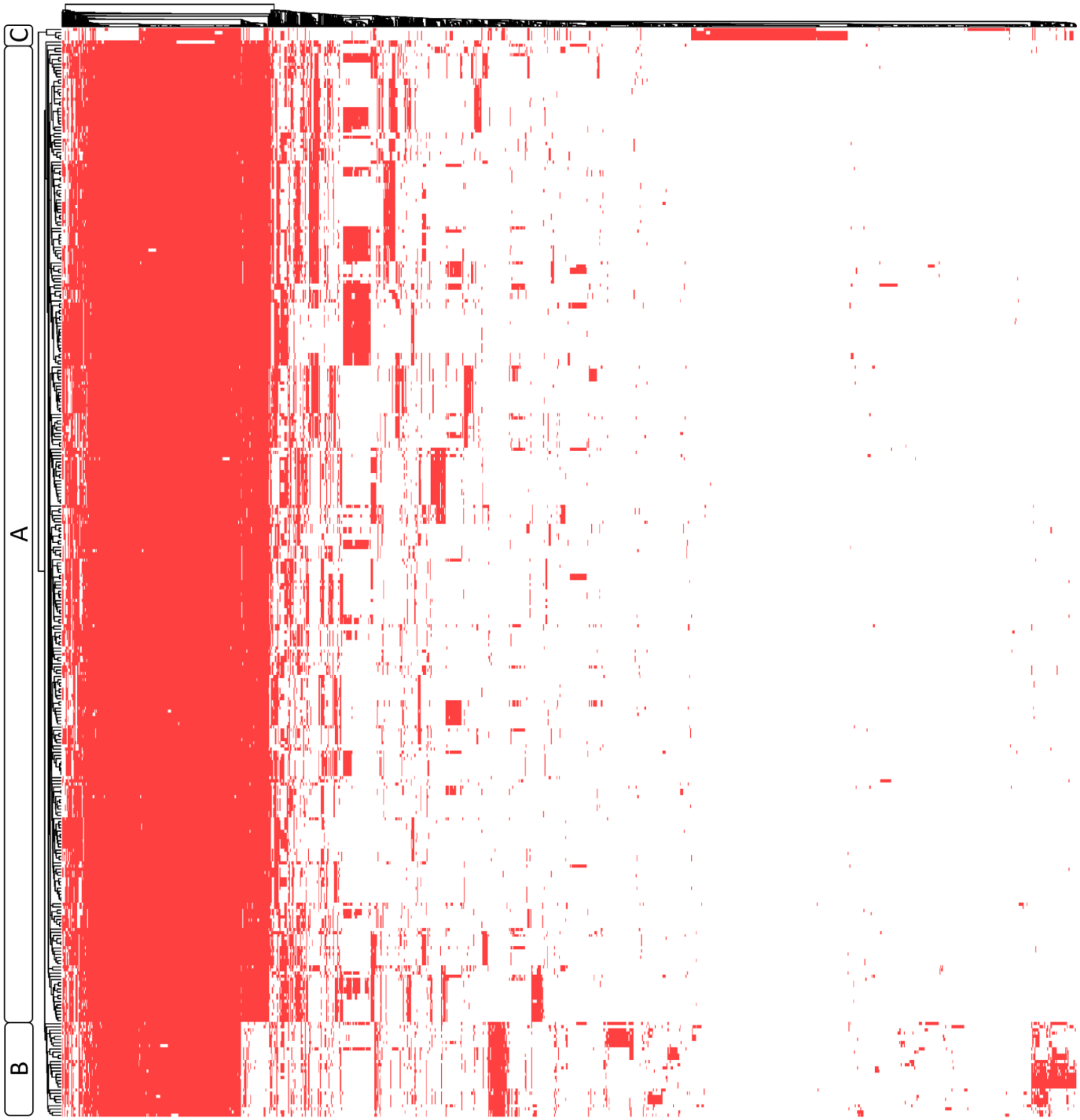


**Figure S1. Accessory genome variability among Phylogroups A, B, and C.** The heatmap shows the presence/absence of accessory genes across the analyzed genomes and is accompanied by a dendrogram clustering the strains based on the similarity of their accessory gene content. Genes present are marked in red. Letters A, B, and C indicate the classification of each strain within the three phylogroups identified through phylogenetic analysis.
