## Supplementary material for "A global perspective on the genomics of Moraxella catarrhalis": Figure S2

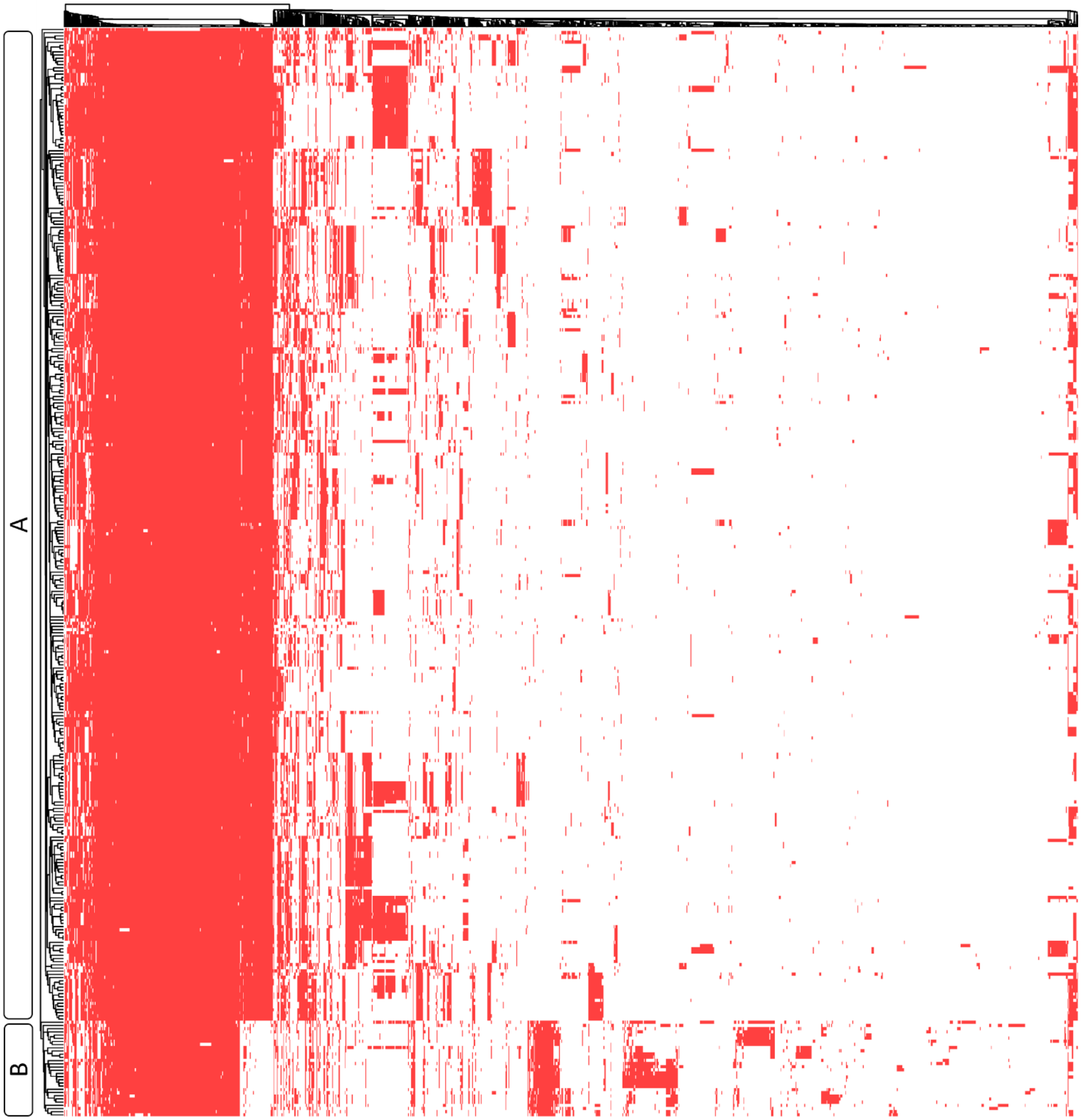


**Figure S2. Accessory genome variability among Phylogroups A and B.** The heatmap shows the presence/absence of accessory genes across the analyzed genomes and is accompanied by a dendrogram clustering the strains based on the similarity of their accessory gene content. Genes present are marked in red. Letters A and B indicate the classification of each strain within the three phylogroups identified through phylogenetic analysis.
